# Hemodynamic Networks Capture Slow Neural Network Persistence but not Faster Behavior-Dependent Network Dynamics

**DOI:** 10.64898/2026.07.05.736600

**Authors:** Lisa Meyer-Baese, Dieter Jaeger, Shella Keilholz

**Affiliations:** Department of Biomedical Engineering, Georgia Institute of Technology and Emory University, Atlanta, Georgia, USA; Department of Biology, Emory University, Atlanta, Georgia, USA

**Author notes:** This work was jointly supervised (Dieter Jaeger, Shella Keilholz).

## Abstract

How can the same spatial pattern of functional connectivity arise from fundamentally different temporal dynamics and biophysical mechanisms? We address this question directly by using simultaneous high-speed wide-field voltage and hemodynamic imaging in awake mice. We demonstrate that canonical functional connectivity networks were preserved across all cortical timescales from infraslow to gamma, yet this shared spatial architecture was generated by completely distinct mechanisms. Infraslow neural and hemodynamic network dynamics were dominated by behavior-dependent state persistence: behavior modulates how long coordinated activity patterns were maintained. In contrast, fast neural network dynamics were shaped by state-dependent selection where behavior determined which spatial patterns are expressed, not how long they lasted. Moreover, low-frequency brain states explained more variance but were less reproducible across sessions, while high-frequency states were more stable and generalizable. These findings reveal that identical functional connectivity can be generated by opposite temporal mechanisms, and establish that hemodynamic signals preferentially reflect the slow, persistent component of neural network activity. Our results provide a mechanistic framework for interpreting fMRI-based connectivity measures and reveal frequency-dependent principles governing how behavior shapes large-scale cortical dynamics.

## INTRODUCTION

Behavior arises from rapid interactions among distributed brain regions that flexibly reconfigure. These coordinated patterns of activity give rise to spatiotemporally structured correlations that enable the brain to route information and perform different cognitive computations at various timescales. Converging evidence suggests that spontaneous changes in behavior and arousal state are tightly linked to transient, recurring whole-brain states in both neural and hemodynamic signals [1, 2]. Yet a fundamental question remains unresolved: can the same spatial patterns of large-scale functional connectivity arise from fundamentally different underlying dynamics, and if so, what does this reveal about how the brain organizes activity across timescales?

There is growing evidence that spontaneous activity can be decomposed into low-dimensional, transient states [3, 4]. This principle generalizes across various signal sources [5–7], from hemodynamic measures, which rely on low-frequency oscillations below 0.1 Hz [8, 9], to electrical activity that fluctuates synchronously across distributed brain regions spanning five orders of magnitude from approximately 0.05 Hz to 500 Hz [10–12]. Additionally, electrophysiology studies to date suggest that a multitude of neural operations can occur in parallel across these timescales [13]. This multiplexing produces multiple dynamic network configurations at different timescales, which in combination are often inferred to represent the slow hemodynamic networks observed [7]. Occurrence of these transient states on the timescale of seconds is known to relate to behavioral and physiological fluctuations [1, 14, 15]. This suggests that there are slow recurring patterns of brain activity that can emerge from or give rise to physiological states that have a stereotyped representation in the brain. Yet it remains unknown whether the mechanism through which behavior shapes these recurring network states is consistent across timescales, or whether the process of behavioral modulation itself varies depending on the frequency band examined.

Prior work examining whether different frequencies support distinct network architectures has largely asked whether different timescales produce different spatial maps of functional connectivity. Here we ask a more mechanistically targeted question: if the same large-scale spatial networks are expressed across multiple frequencies, do they arise through the same mechanisms? And does behavior reshape these shared networks through the same process at every timescale, or through fundamentally different ones?

To address these questions, we used simultaneous high-frame-rate wide-field voltage imaging and hemodynamic imaging in awake mice, enabling direct comparison of neuronal and vascular signals across behavioral conditions. We simultaneously tracked spontaneous changes in behavioral state defined by changes in orofacial movements and fidgeting. We show that canonical resting-state networks were preserved across all neuronal frequency bands from infraslow to gamma, establishing a shared spatial scaffold that was largely frequency-invariant. Despite this, the temporal dynamics underlying these networks diverged substantially across timescales. Low-frequency and hemodynamic activity were dominated by temporally persistent network states whose expression was strongly linked to prolonged periods of spontaneous behavior. Whereas higher-frequency activity was characterized by rapid state transitions in which behavior shapes which network configurations were recruited rather than how long they were maintained. Furthermore, low-frequency brain states explained more variance within a session but were less reproducible across sessions, while higher-frequency states were more stable and generalizable. Hemodynamic dynamics most closely paralleled infraslow neural activity in both their temporal structure and their mode of behavioral modulation, providing a mechanistic account of what fMRI resting-state signals primarily reflect. Together, these findings demonstrate that identical functional connectivity can be generated by contrasting mechanisms and reveal frequency-dependent principles through which behavior shapes the persistence and expression of large-scale cortical network dynamics.

## RESULTS

### Simultaneous High-Speed Neural and Hemodynamic Imaging Captures Cortical Network Activity Across Timescales

Neural and hemodynamic activity across the dorsal cortex was recorded simultaneously using wide-field optical imaging in five awake, head-fixed mice. These mice expressed a voltage-sensitive fluorescent protein, JEDI-1P-Kv2.1, which restricts membrane protein expression to the soma and proximal dendrites. A Cre-dependent adeno-associated JEDI-1P vector was injected into the ventricles of EMX1-Cre mice at p0 to obtain preferential expression in excitatory neurons across the cortex [16]. To correct for hemodynamic absorption, these mice also expressed a red reference fluorescence, mCherry (AAV9-hSyn-mCherry). Both fluorescent molecules can be excited with a single blue LED light source and simultaneously measured after splitting the emission signals using two CMOS cameras (**Fig1A**). An additional red LED was used on a subset of trials to record intrinsic red reflectance as an alternative measure of hemodynamics. To allow analysis of a broad range of frequencies, trials were recorded with a frame rate of 200 frames per second. Each recording session consisted of whisker stimulation (30Hz, air-puff stimulation delivered for 3s to one side of the face per trial) and resting state trials (**Supp Fig1-1A,B**). Each mouse underwent several imaging sessions across days, resulting in 40 minutes’ worth of resting-state data per mouse.

**Figure 1.**
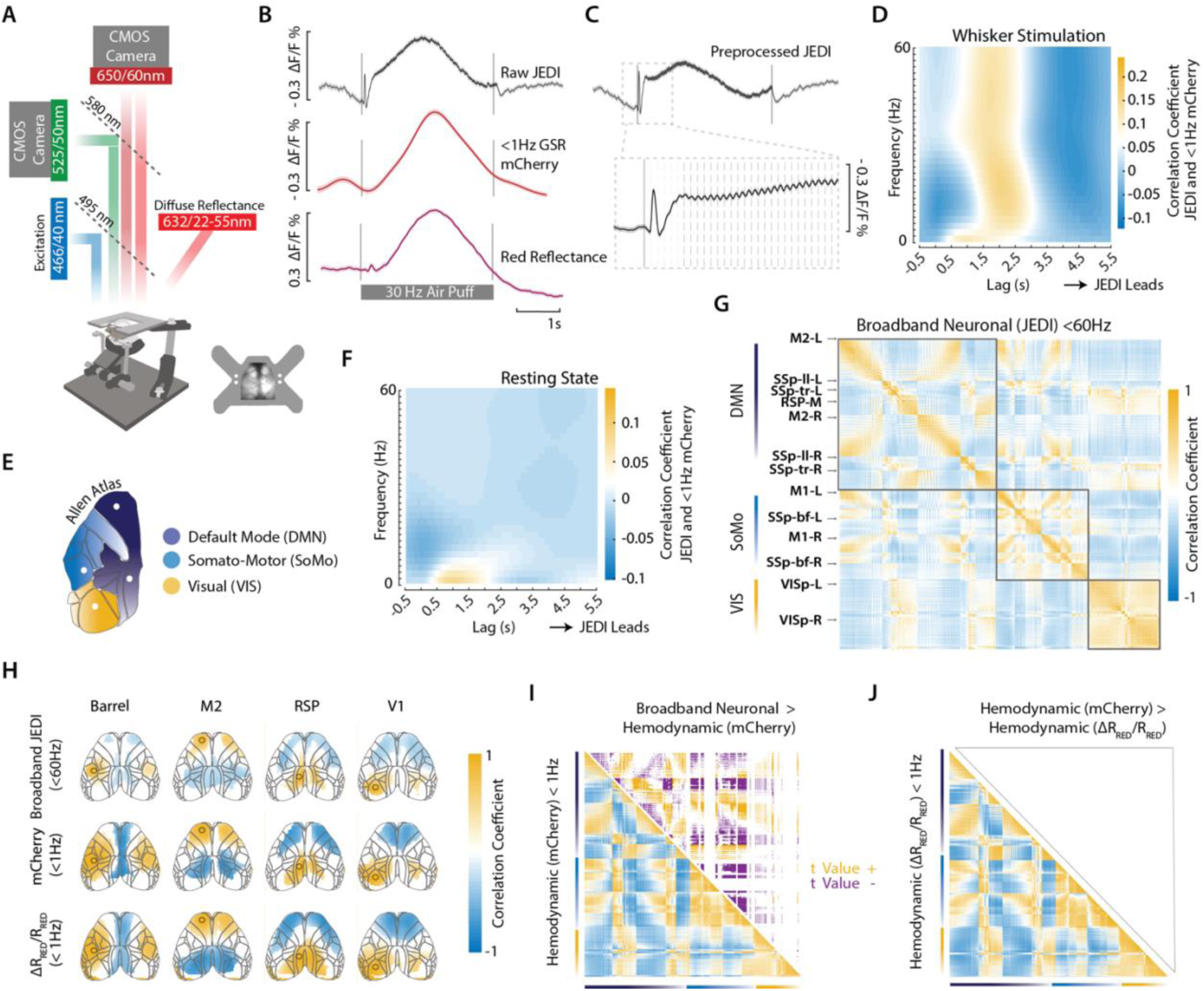
Wide-Field Optical Imaging of Voltage Membrane Potentials and Hemodynamics. **A)** Schematic of wide-field imaging setup consisting of two different excitation LEDs and two CMOS cameras that can simultaneously acquire up to 200 fps. **B)** Top trace (black) shows the raw trial average JEDI-1P-Kv trace from barrel cortex for a 3-second-long, 30Hz air puff stimulus. This trace is before any hemodynamic regression. Middle trace (dark red) shows the simultaneously acquired <1Hz GSR mCherry trace. Bottom trace (purple) shows the trial average <1Hz red reflectance data obtained from the same mice. For top two traces trial average represents n = 540 trials from 5 mice. For the bottom trace data reflects trial average of n = 562 trials from 4 mice. Shaded error bars in all plots denote the standard error. Grey lines denote the start and end of the whisker stimulation **C)** Same data as in panel B showing the trial average JEDI-1P-Kv trace following the sequential hemodynamic regression. Inset shows the initial fast voltage response and the 30Hz oscillation. **D)** Displayed are the cross-correlation values obtained from barrel cortex between band-pass filtered versions of the JEDI-1P-Kv signal and the <1Hz GSR mCherry signal for whisker stimulation data shown in panel C. The frequency resolution on the y-axis is 2Hz. **E)** Schematic illustration of the 3 network parcellations used for constructing the FC matrix. **F)** Same as Panel D but for activity in left and right barrel cortex for the resting state conditions representing n = 540 trials. **G)** Average static FC matrix for broadband <60Hz GSR preprocessed JEDI-1P-Kv signal across all areas in the dorsal cortex n = 289 trials. **H)** Average resting state seed-based FC maps for four different seeds/ROIs represented by each column. Each ROI is between 26-28 pixels in size. Top row represents the broadband <60Hz GSR preprocessed JEDI-1P-Kv signal. Second row is the simultaneously acquired <1Hz GSR mCherry signal. The bottom row is the <1Hz GSR red reflectance data. Top two rows represent trial average data from n = 289 trials bottom row is from n = 120 trials. All correlation values are plotted using the same colorbar limits set to range from -1 – 1. Pixels are thresholded to pixels with significant connectivity as determined using a two-sided t test corrected for multiple comparisons with false discovery rate (FDR). **I)** Bottom left is the low-passed < 1 Hz GSR mCherry signal FC matrix. The top right is the two-sample two-sided t-Test results comparing broadband static FC voltage to hemodynamic activity. Corrected for multiple comparisons with FDR. Displayed here are the t values for matrix entries that have statistically significant correlation values. They are thresholded to represent positive and negative correlations. **J)** Bottom left is the GSR red reflectance data FC matrix. The top right is the two-sample two-sided t-Test results comparing the low-pass mCherry signal to the red-reflectance data hemodynamic activity using the same statistics as in panel I. No statistically significant differences were found.

**Supplemental Figure 1-1.**
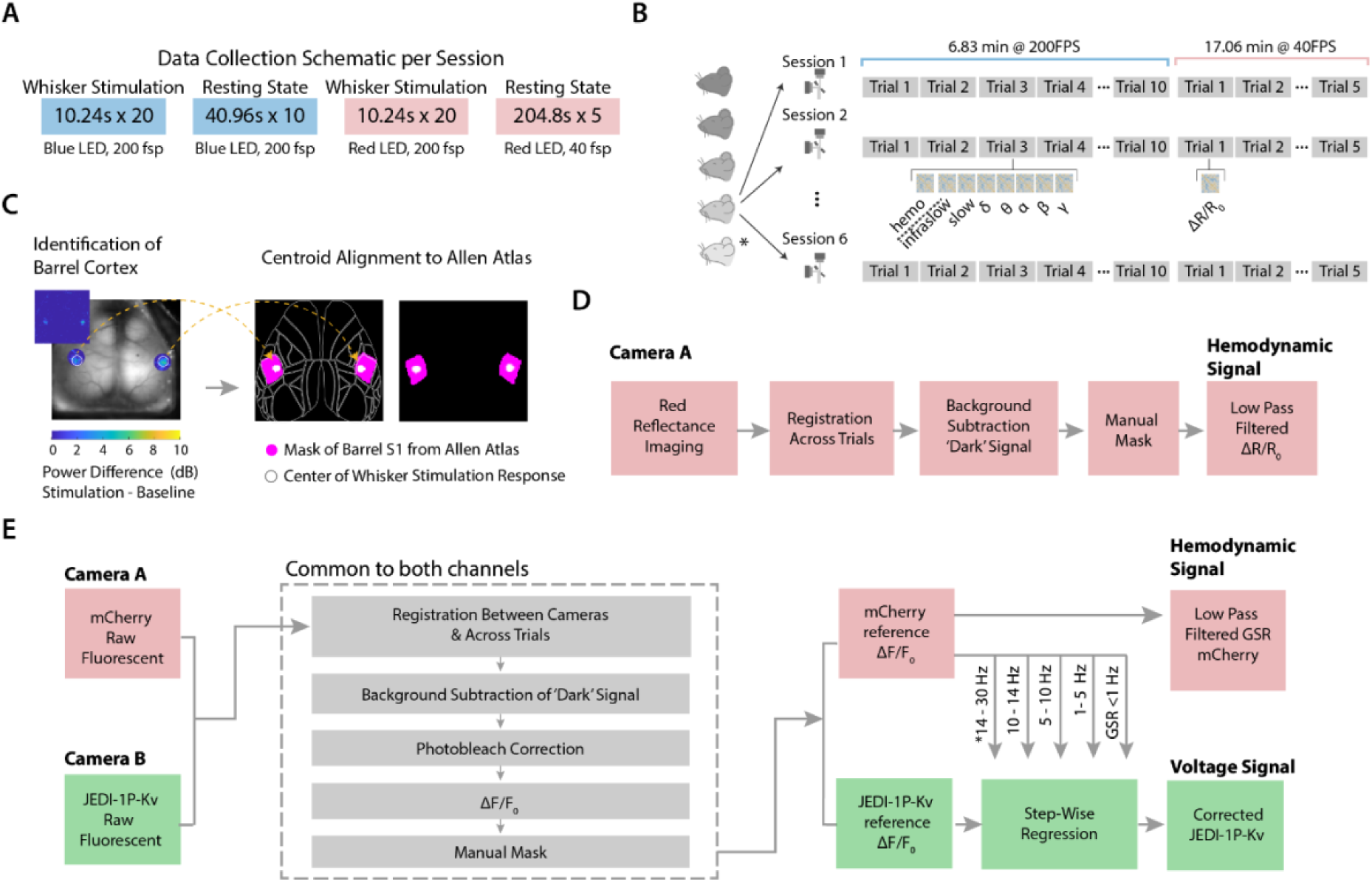
Data Collection and Preprocessing. **A)** Schematic illustrating the different trial types recorded during one imaging session. Top row denotes the trial type which is either a whisker stimulation trial or resting state trial. Directly underneath in the colored boxes are the durations and number of trials recorded for each type. The color of the block denotes which LED was used. Underneath each colored block there is labeled the LED type and the frame rate used for that block of trials. **B)** The schematic extends on what is shown in panel A to highlight group-level data. Data from four mice consist of 6 resting state trials with both blue and red LEDs. For one mouse there is only blue LED data across 3 sessions that were twice as long, resulting in the same amount of net resting state data. **C)** Cross-session alignment to the Allen Atlas was performed using the whisker stimulation data. An image showing the power difference between stimulation and baseline was obtained for each whisker stimulation session. Pixels with increased power represent the whisker barrels (left), which define the two ROIs. Using centroid alignment these two ROIs are then co-registered to the centroid of a mask of barrel cortex from the Allen Atlas (right). **D)** Preprocessing steps applied to the red reflectance imaging from a single channel **E)** Preprocessing pipeline for the dual camera image acquisition. Steps common to signals from both cameras are in gray boxes. Hemodynamic correction is applied using band-pass filtered versions of the mCherry signal that are regressed out of the JEDI-1P-Kv signal. The corrected JEDI-1P-Kv signal represents the voltage signal referenced throughout the paper. The simultaneously acquired hemodynamic signal refers to the low pass filtered global signal regressed mCherry signal.

A red reference channel for hemodynamic correction has become common in several voltage imaging studies that are interested in fast neural signaling [16–18]. This method effectively removes artifacts across a range of frequencies through a sequential regression preprocessing step, which has been validated using JEDI-1P in our lab (**Supp Fig1-1E)** [16]. More importantly, this method eliminates the need for interleaving additional LED sources for diffuse reflection imaging, which would limit the maximum sampling rate of each color channel. In the present study, we leveraged the red reference channel not only for hemodynamic correction but also as a proxy for hemodynamic signaling. To this end, we show through modeling and experimental data that the slow global signal regressed (GSR) <1Hz mCherry signal mirrors the expected hemodynamic response one would observe with more traditional optical hemodynamic measures like diffuse reflectance imaging. Unlike traditional diffuse reflectance imaging, the measured signal from a red reference channel is a composite of both the excitation and emission wavelengths and their respective absorption of oxygenated (HbO) and deoxygenated (HbR) hemoglobin (**Supp Fig1-2F)**. Despite being a composite signal, the mCherry fluorescence signal is primarily weighted towards changes in oxygenated hemoglobin (HbO) (**Supp Fig1-2G)**. This can be observed in the shared slow hemodynamic transient that is present in the uncorrected JEDI signal (**Fig1B).** The time-lagged relationship between the corrected green voltage fluorescence and the slow <1 Hz mCherry hemodynamic signal for a 3-second whisker air puff was assessed by looking at the cross-correlation between filtered versions of the JEDI signal and the slow hemodynamic signal (**Fig1D**). This revealed a physiologically meaningful time lag (peak correlation of 0.24 centered at 1.5 s). The JEDI stimulus transient (Fig1C) extends beyond the slow frequencies and can be faithfully observed in the cross-correlation up to 60 Hz (**Fig1D**). For resting-state trials, this same time-lagged correlation structure is observed in barrel cortex, with a lower peak correlation of 0.15 that drops off at 8Hz (**Fig1F**). This mirrors results seen using the LFP signal as a measure of neuronal activity [19].

**Supplemental Figure 1-2.**
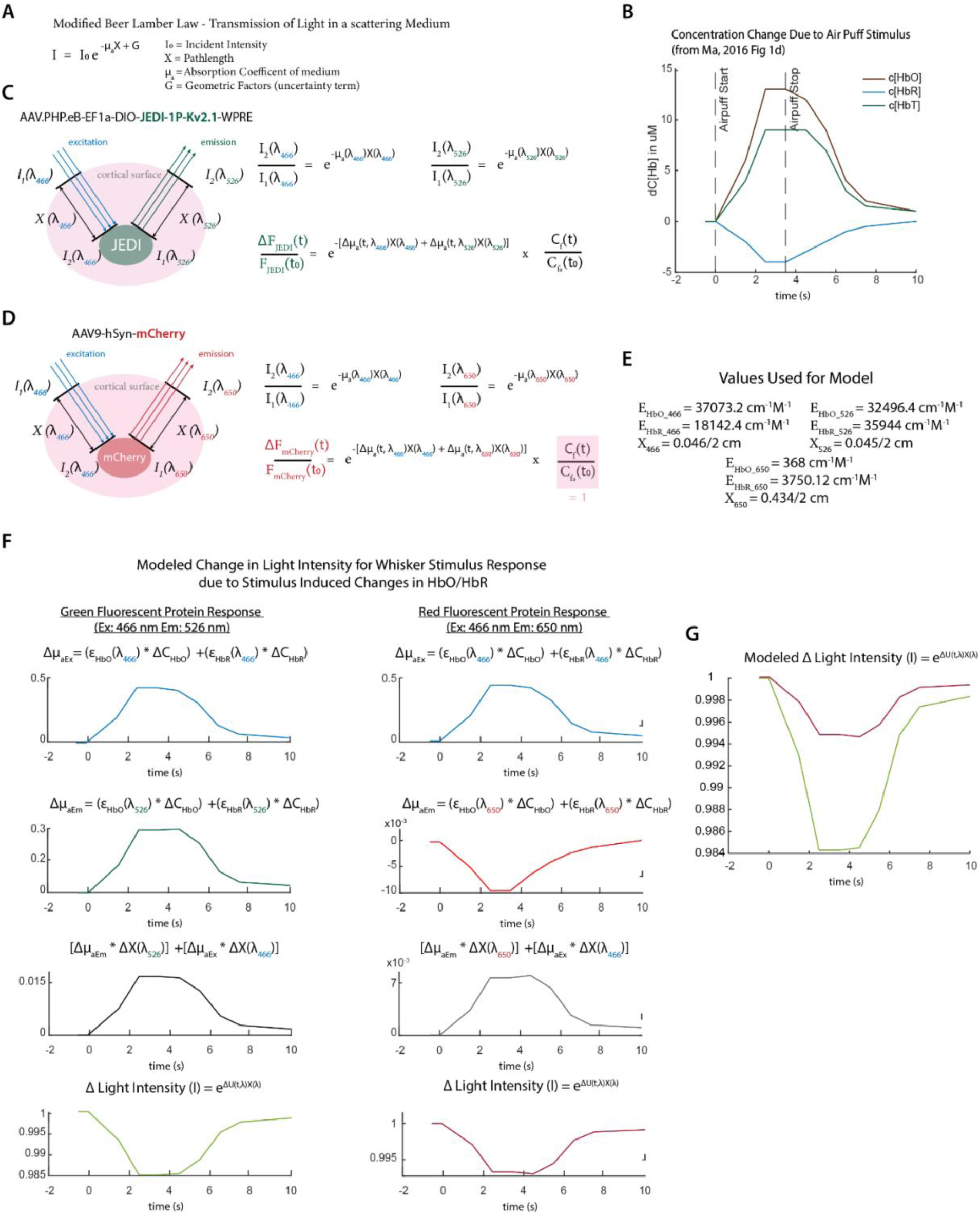
Modeled Hemodynamic Contributions to Fluorescent Signals. **A)** Light traveling through a scattering medium like the brain can be modeled through the Modified Beer’s Law. **B)** Changes in concentration in barrel cortex of HbO (brown), HbR (light blue) and HbT (green) for a whisker stimulus as reported in [20]. **C)** Schematic of variables to consider for fluorescent imaging of a GFP such as JEDI-1P-Kv (left) and mathematical formulation (right) to describe change in measured light intensity based on a formula in panel A. **D)** Same as in panel C but for a RFP like mCherry. **E)** Modeled values from the literature for the extinction coefficients for HbO and HbR, along with the reported values for optical pathlength for all 3 wavelengths of interest [20]. **F)** Left column represents the modeled change in light intensity for JEDI-1P-Kv using the values using the formula in panel C and the values from panel B and E. From top to bottom are the modeled changes in light for: excitation wavelength, emission wavelength, both combined with factoring in the pathlength traveled, the final measured change in light intensity. Right column represents the same thing but for mCherry. Note the difference in magnitude on the y-axis for the excitation and emission wavelengths of mCherry. There is a stronger absorption due to HbO and HbT at the excitation wavelength. **G)** An overlay of the modeled change in light intensity due to neurovascular coupling that would be observed for both JEDI-1P-Kv (green) and mCherry (red) during the whisker stimulus response. For both fluorescent signals a negative change in fluorescence due to neurovascular coupling is expected.

The spatial organization of resting state networks was probed using seed-based correlation mapping. Connectivity maps for the broadband JEDI-1P signal reveal the expected interhemispheric connectivity (FC) maps for several different ROIs across the dorsal cortex (**Fig1H, top row**). Similar maps can be obtained from the <1Hz mCherry signal, which also reflects bilateral resting-state networks that are less spatially localized than the neuronal counterparts (**Fig. 1H, middle row**). An important control for all these measures is the alternative hemodynamic signal representing diffuse reflectance of red light, which was acquired in a subset of mice (n = 4) (**Supp Fig1-1B and Fig1-1D**). The observed hemodynamic response in the red reflectance mirrors the slow temporal kinetics of the response seen in the mCherry and in the raw JEDI-1P signal (**Fig1B**). During resting state, the observed static FC maps for the red reflectance are spatially similar to those obtained from the mCherry (**Fig1H, bottom row**).

**Figure 2.**
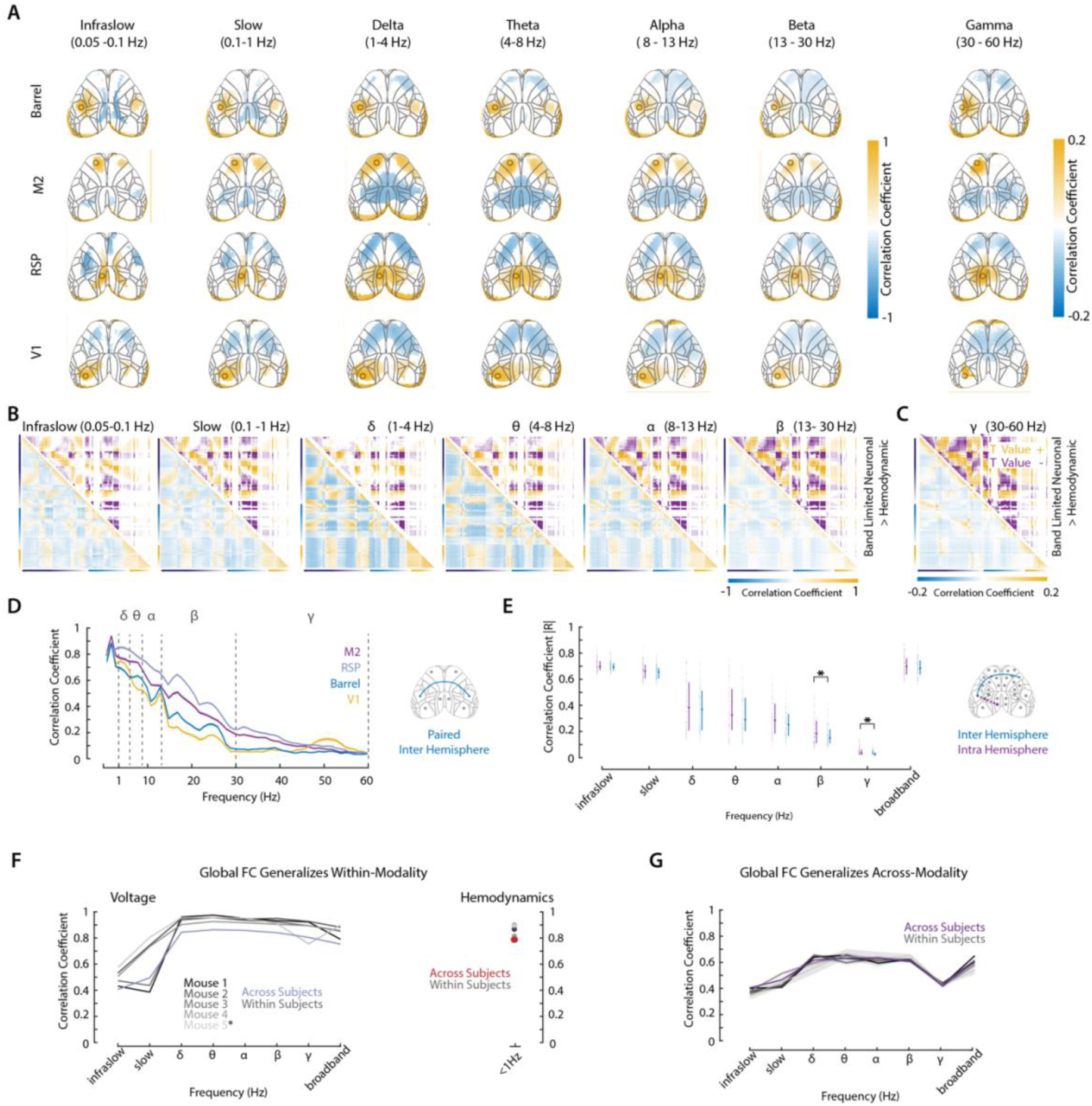
Preserved and Reliable Neuronal Bandlimited Functional Connectivity Across Canonical Frequency Bands. **A)** Average static seed-based FC for bandlimited neuronal signal filtered to represent canonical frequency bands. Pixels are thresholded to pixels with significant connectivity as determined using a two-sided t test corrected for multiple comparisons with FDR. **B)** Bottom left half represents the average static FC matrix for the bandlimited neuronal signals. Top right is the two-sample t-test results comparing the bandlimited neuronal FC to that from the hemodynamic signal. Thresholded t values are shown to represent significant positive and negative values. Corrected for multiple comparisons using FDR. **C)** Same as in B but for the gamma bandlimited signal whose FC matrix is plotted using a different color bar axis limit. **D)** Interhemispheric correlation for 4 different regions similar to the ones used for the static FC connectivity matrices, except that these represent only a single 1 x 1 pixel from each hemisphere rather than a larger circular ROI. The frequency resolution on the x-axis is 2Hz. Shaded bars represent SEM. **E)** The absolute value of the correlation obtained from 16 1×1 pixel ROIs spread out across dorsal cortex (schematic on the right, grey dots) from the left hemisphere correlated with all possible pairs of points on the right hemisphere. (16 choose 2 = 120 unique pairs for interhemispheric connections in blue) and the same is done for all possible pairs within a hemisphere (interhemispheric correlations in purple). Each dot represents the correlation value from one pair of points. The overlaid boxplots represent the group-level median (n = 289 trials across 5 mice), along with the first and third quartile. Statistical significance between inter- and intra-hemispheric connectivity for each frequency band was determined using a Mann-Whitney U test with FDR correction. Significance thresholds with asterisks denote p < 0.05 *, p<0.01 **, and p < 0.001 ***. **F)** Within-modality correlation for pairs of FC matrices. On the left is the correlation coefficient for all pairs of FC matrices derived from the band-limited JEDI signal, either within a subject (grey lines) or across subjects (light purple line). Error bars represent SEM. On the right is the same but for the <1Hz mCherry FC matrices within subjects (grey dots) and across subjects (red dot). **G)** Across modality correlation between JEDI and mCherry FC matrices within (grey) and across subjects (purple). Error bars represent SEM.

The seed-based FC results were corroborated by the pixel-wise mapping of interareal connectivity via static FC correlation matrices. Recorded activity from our field of view (FOV) spans 3 canonical resting state networks: the default mode (DMN), the somatomotor (SoMO) and the visual network (VIS) (**Fig1E**). To help anatomical interpretability, pixels in the FC matrices were assigned to one of the 3 networks (**Fig1G, Fig1I and Fig1J**). The static FC matrix for the broadband JEDI signal highlights high within-area connectivity and interhemispheric correlations across dorsal cortical regions. It also showcases the within hemisphere anticorrelation between anterior and posterior regions (**Fig1G**). These results were also present in the static mCherry FC and red reflectance matrices while being spatially more blurred (**Fig1I and Fig1J**). Comparison of the FC mCherry matrices to the voltage matrices reveals no statistically significant differences within the VIS network. For the SoMo network mCherry signal is significantly more positively correlated within a given region compared with the JEDI-1P signal (**Fig1I, top right**). When comparing the mCherry FC matrices to the red reflectance FC matrices no statistically significant differences appear (**Fig1J, top right**).

This method establishes a platform for directly comparing neuronal and hemodynamic network dynamics across timescales. Although neural and vascular signals exhibit distinct temporal properties, both reveal a common large-scale organization of cortical networks. This shared spatial architecture, measured simultaneously within the same animals, provides a foundation for investigating how both neuronal activity across frequencies and hemodynamic signals underpin functional connectivity measurements.

### A Common Spatial Architecture Underlies Cortical Networks Across All Frequency Bands

The preservation of large-scale network organization across neuronal and hemodynamic signals raises the question of whether a similar organizational principle extends across neuronal timescales. If functional connectivity reflects a fundamental property of cortical organization, canonical network structure should remain evident despite large differences in the temporal characteristics of the underlying activity. To test this, we broke down the recorded JEDI-1P signal into canonical frequency bands: infraslow [0.05-0.1Hz], slow [0.1-1Hz], delta [1-4Hz], theta [4-8Hz], alpha [8-13Hz], beta [13-30Hz], and gamma [30–60]. Using the filtered, band-limited JEDI-1P signal we calculated seed-based FC maps and FC matrices (**Fig2A,B, and C**). This analysis revealed a conserved network organization across all frequency bands for all four selected seeds: Barrel, M2, RSP and V1 seeds, with all 4 seeds exhibiting significant correlations and anticorrelations following multiple comparison corrections (**Fig2A)**. This preserved connectome structure was corroborated by the static FC matrices (**Fig2B** and **Fig2C,** bottom left triangle). In contrast, the mCherry signal exhibited this canonical network structure, which included the anticorrelation, only in the low frequencies up to the delta band (**Supp Fig2-1 B)**. Statistically significant anticorrelations in the static FC analysis did not extend past the slow frequency (**Supp Fig2-1 A)**. Higher-frequency pixels were broadly correlated with one another, likely due to a broad distribution of common non-neuronal sources such as respiration and heart rate that are present in the mCherry signal (**Supp Fig2-1 A-B**). When comparing the band-limited JEDI FC matrices to the <1Hz mCherry matrix, there is an increasing number of matrix elements that differed between the two as a function of increasing frequency (**Fig2B**, top right triangle). This was assessed using a t-test comparing z-transformed bandlimited JEDI FC correlation values to hemodynamic FC correlation values. Plotted are the positive and negative thresholded T values corrected for multiple comparisons. On average the hemodynamic signal has higher connectivity for pixels within the somatosensory network compared with the band-limited slow–gamma band JEDI signal (**Fig2B**, top right triangle, purple T < 0). The JEDI signal across frequencies exhibits significantly stronger correlations within the DMN and with the DMN and pixels from SoMo and VIS networks (**Fig2B**, top right triangle, yellow T > 0).

**Supplemental Figure 2-1.**
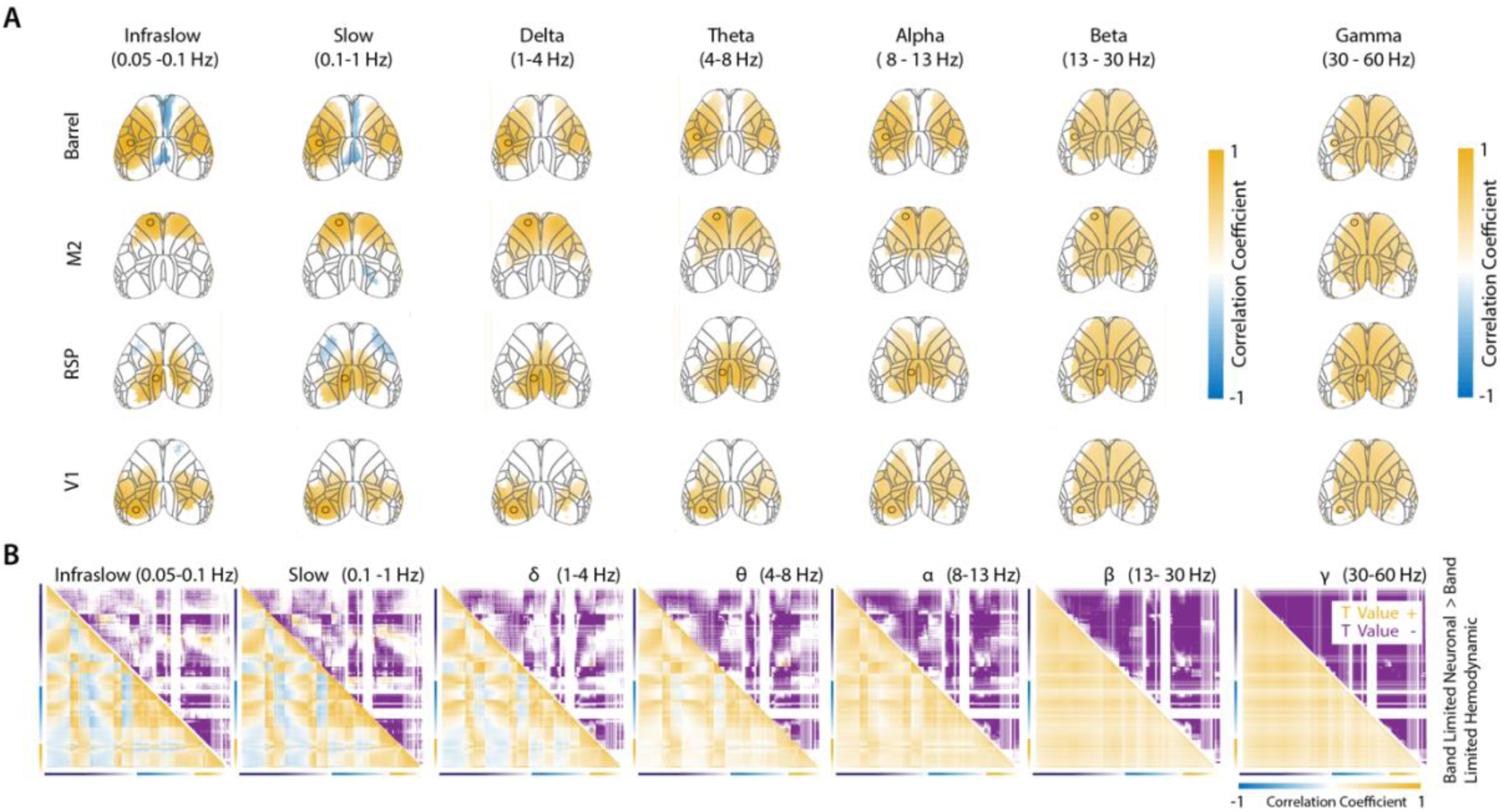
Different Bandlimited Functional Connectivity Across Faster Frequency Bands in mCherry Signal. **A)** Average static seed-based FC for bandlimited mCherry signal processed and filtered in the same way as the corrected JEDI signal. Pixels are thresholded to pixels with significant connectivity as determined using a two-sided t test corrected for multiple comparisons with FDR. **B)** Bottom left half represents the average static FC matrix for the bandlimited mCherry signals. Top right is the two-sample t-test results comparing the bandlimited neuronal FC to that from the bandlimited mCherry FC. Thresholded t values are shown. Corrected for multiple comparisons using FDR.

The degree of interhemispheric voltage correlation between paired regions (i.e. left and right barrel cortex) ranged from 0.7 to 0.9 and decreased as a function of increasing frequency for various areas in cortex (**Fig2D**). This held not only for paired regions but also for general connectivity within and across hemispheres (**Fig2E**). Despite this drop-off, several areas remained significantly correlated (**Fig2A)** through the gamma band. While both the inter- and intrahemispheric connectivity decreased with increasing frequency, the degree of intrahemispheric connectivity remained significantly higher than the interhemispheric connectivity for the higher frequencies (**Fig2E**).

We evaluated the reliability of the global topography structure as represented through the static FC matrices within and across modalities by comparing the correlation of the average FC matrices across sessions. Within the JEDI signal the within-subject FC matrices were consistently highly correlated across delta – gamma band (≥ 0.8), with infraslow and slow matrices having the lowest correlation in terms of FC structure across sessions. Within each subject, the correlation structure is also more consistent on average across sessions (grey lines, **Fig2F**) than across mice (purple line, **Fig2F**). The same is true for the FC matrices derived from the mCherry signal (**Fig2F** grey dots for within mice, red dot for across mice). Global across-modality structure between JEDI and mCherry was also high but lower than the within-modality correlations. For delta–beta and broadband the correlation was ≥ 0.6 while for infraslow, slow and gamma it was ≥ 0.4 (**Fig2G**). Additionally, minimal differences were observed within (grey lines) versus between subjects (purple line, **Fig2G**).

Together, these findings demonstrate that large-scale cortical functional networks are preserved across a broad range of neuronal timescales. Although the magnitude of connectivity progressively decreased with increasing frequency, the overall spatial organization of canonical networks remained remarkably stable. The high reproducibility of this network structure across sessions, animals, and imaging modalities further suggests that static functional connectivity captures a robust and conserved feature of cortical organization. Importantly, this frequency-invariant spatial scaffold provides the interpretive foundation for the dynamic analyses that follow: because the same large-scale networks are present across all timescales, differences in how those networks are expressed over time cannot be attributed to differences in spatial architecture, but must instead reflect differences in the temporal dynamics in which a common network repertoire is organized and expressed. This motivated a closer examination of how network dynamics unfold across frequencies and behavioral states.

### Low-Dimensional Dynamic Functional Connectivity Unfolds Concurrently at Different Timescales

So far, we have analyzed this resting state data within the context of static FC, which looks at time-averaged signal correlations across areas. However, resting state activity is known to be highly dynamic, resulting in several different network configurations that can unfold throughout the duration of a single resting state session [3, 4]. To capture these resting state dynamics, we extracted co-activation patterns (CAPs), which are commonly used in dynamic rsfMRI analysis [21]. Similar to other work that uses CAPs on preclinical neuroimaging data [22], we ran this analysis without any *a priori* intensity-based restrictions. The CAPs obtained represent recurring, momentary network-level activity patterns that characterize different brain regions that are concurrently activated or deactivated. CAPs are also defined at the temporal resolution of the data, making them well-suited for this cross-frequency study.

CAPs were identified on a per-session basis for all signals and frequency bands that we also used for the static analysis. This includes the slow mCherry hemodynamic signal, the broadband JEDI voltage signal, along with the band-limited JEDI signals (**Fig3A**). CAPs were obtained by running k-means clustering on concatenated z-scored data on a per-session and per-frequency basis for clusters ranging from 2-26. This results in K number of CAPs based on the selected K cluster number. Each CAP is an image obtained from averaging all frames together that were clustered based on their spatial similarity. The optimal number of CAPs for each signal type was then obtained by looking at the elbow point in the gain of percent variance explained as a function of increasing K (**Supp Fig3-1A)**. The mean cluster number K, and therefore the mean number of CAPs did not greatly differ across signal types (**Fig3B, purple dots**) ranging on average between 13 (for theta) to 20 (for hemodynamics, delta, gamma and broadband signals) CAPs. Despite similar number of ideal CAPs, what did vary across signal types was the percent of the variance that these CAPs explained in the data. Time-varying changes in hemodynamic resting state activity were best explained by CAPs, accounting for an average of 65% of the explained variance (**Fig3B, teal dots**). This then steadily decreased with increasing frequency in the voltage signal which starts at 46% for infraslow and drops to 5% for gamma band (**Fig3B, teal dots**).

**Supplemental Figure 3-1.**
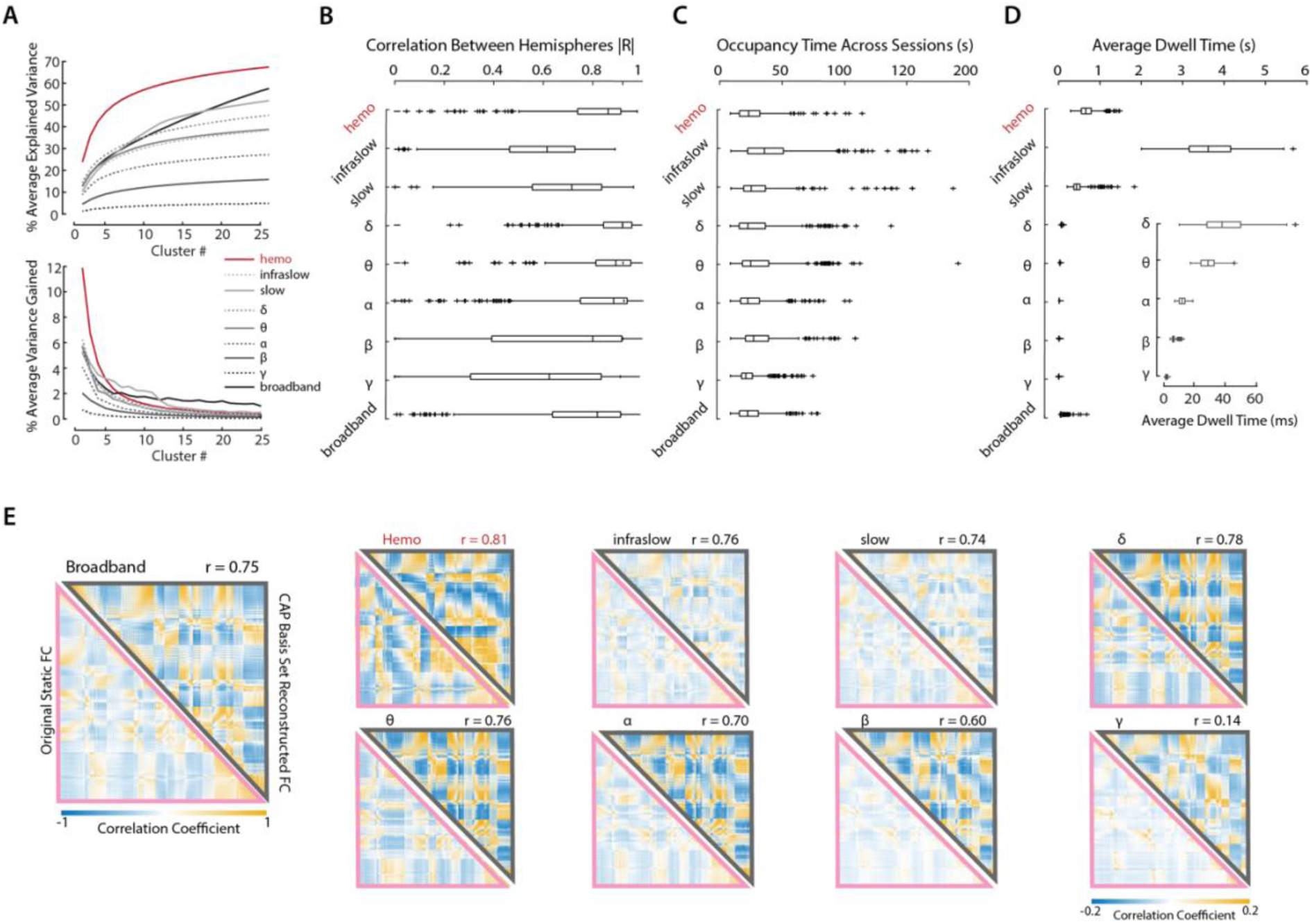
CAP Properties Across Frequency Bands. **A)** Session average change in average explained variance and variance gained as a function of increasing cluster number K across different signal types and frequencies. **B)** Distribution of the absolute value of the correlation between left and right hemispheres. Sorted as a function of frequency bands. **C)** Distribution of the average percentage of all imaging frames assigned to each CAP as a function of different frequencies. **D)** Distribution of the dwell time in seconds, which represents how much time is spent in a given CAP state, plotted as a function of different frequencies. The bottom excerpt shows the same dwell time in units of ms for theta through gamma band frequencies. **E)** Bottom left pink triangle represents the average FC matrix obtained from the raw data. This is the same data as in Figure 1G shown here to highlight a comparison to the top triangle. Top grey triangles represent the average matrix obtained from reconstructing session-level data based on individually selected ideal k CAPs. The spatial correlation between the two matrices is displayed to the right of each square. Represented is the data for all frequency bands and signal types.

**Supplemental Figure 4-1.**
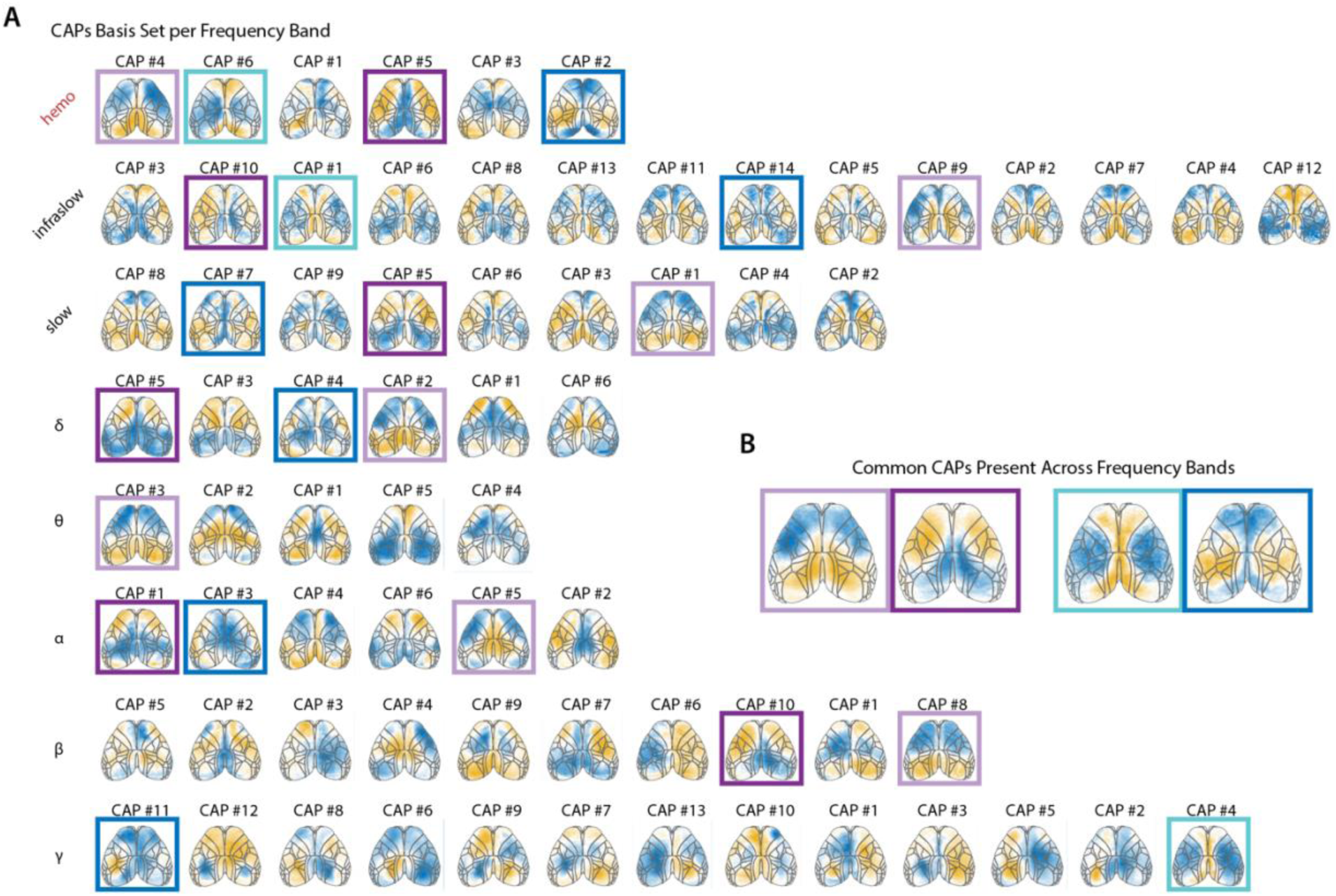
CAP Basis Set Across Frequencies. **A)** The basis set obtained from clustering all the single session frequency specific CAPs with a cutoff of 0.8. Colors underneath certain CAPs highlight repeated spatial patterns that are present across frequency bands **B)** Images obtained from averaging the CAPs highlighted with the respective colors. These images represent the average spatial patterns that are present across frequencies. These 4 average images are grouped into two anticorrelated pairs.

**Supplemental Figure 5-1.**
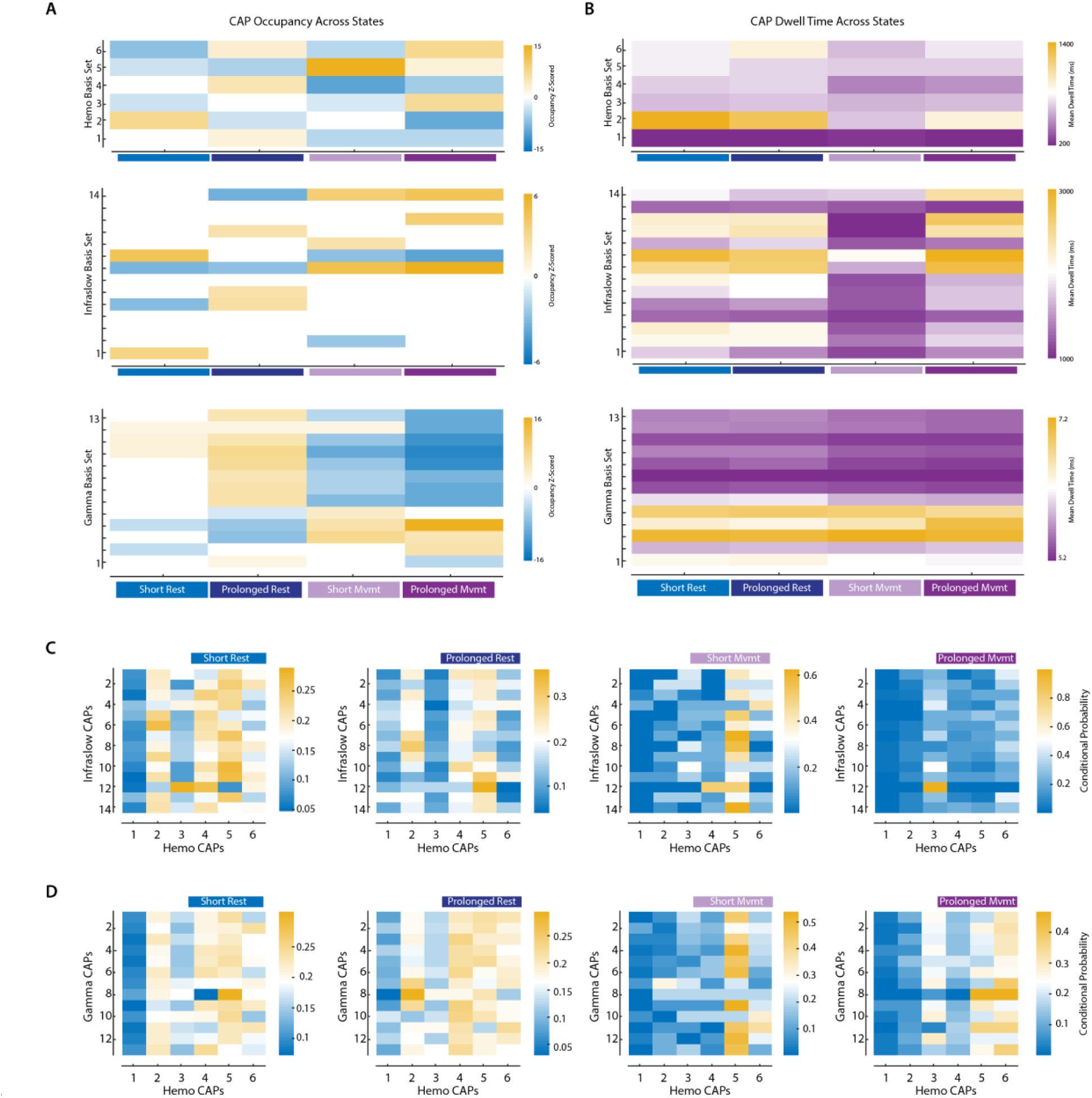
CAP Occupancy and Dwell Time Across States. **A)** Same as Figure 5D, but for the three other signals that were consistent across CAP cutoffs. **B)** Same as Figure 5E but for the three other signals that were consistent across CAP cutoffs.

Figure 3C shows an example of six CAPs from a single session, sorted by dwell time across signal types. A defining characteristic of CAPs across signal types and frequency bands is their spatial configuration which represents an assembly of regional substrates that encompass previously characterized wide-field resting state networks. An example is the motor resting state network which corresponds to bilateral activation of primary and secondary motor cortex, that is present in CAPs #18 and #6 for slow and delta bands, respectively. Several CAPs within a frequency band also configure into state and anti-state pairs characterized by opposing patterns of functional co-activation, such as CAP #3 and #6 for delta band (**Fig3C**). Most CAPs for slower frequencies are symmetric across the midline. However, for beta and gamma band CAPs start to represent more spatially localized patterns of activity that are not necessarily symmetric across the midline. This can be represented by quantifying the absolute value of the correlation between left and right hemispheres in each CAP, which on average goes down and becomes more broadly distributed for frequencies from the alpha-gamma band (**Supp Fig3-1B)**.

**Figure 3.**
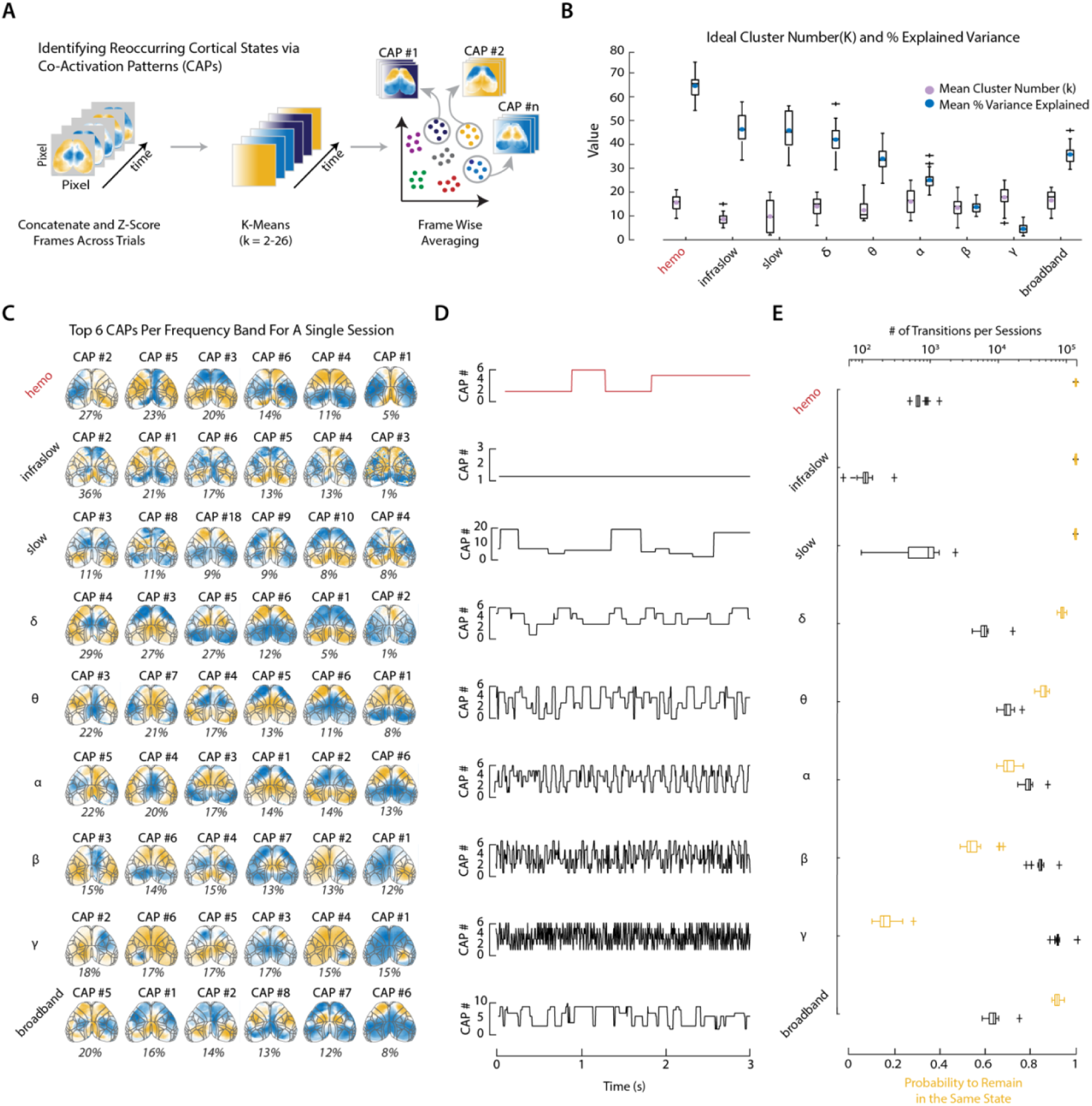
Low-dimensional Spatial Patterns Capture Resting-State Activity that Concurrently Unfolds at Different Timescales. **A)** Schematic illustrating how session-specific CAPs are obtained for each frequency band. K-means clustering is run on a per-session and per-frequency basis for cluster values from 2 – 26. **B)** Purple circles represent the mean K value for each of the frequency bands. The histogram represents the spread of all “ideal” K values selected for each session. This K value was obtained by identifying the elbow point in the explained variance gained. The teal circles represent the average % variance that is explained by the selected K values. The histograms represent this distribution across sessions. **C)** A single session example of the top six CAPs as sorted by % dwell time that are obtained for each frequency band. **D)** Represents the time-varying changes in CAPs. Each row corresponds to the activity patterns for each frequency band in panel C. Data is shown for a 3s window of time. **E)** Grey histograms represent the number of transitions between CAPs for all sessions as a function of different frequency bands. Transition number is plotted on a log scale. Yellow histograms represent probability of remaining in a given CAP state as a function of different frequency bands.

Each original frame can be assigned to a given CAP (**Fig3D**) to then assess the time-varying dynamics. The number of transitions increases with increasing frequency (**Fig3E, black boxplots**), which in turn results in a decreased probability, and therefore a decrease in the average dwell time that is spent in the same state as a function of increasing frequency (**Fig3E, yellow boxplots, Supp Fig3-1D)**. Due to the similar number of CAPs across signal types, the average time that is spent in a given CAP is relatively similarly distributed across signal types (**Supp Fig3-1C)**.

The obtained single-session CAPs reveal dynamic brain states that provide the building blocks of overall brain connectivity, which is assessed with static networks, as is reflected by the high degree of spatial similarity between the original static FC matrix and that obtained by reconstructing the data using CAPs (**Figure 3-1E)**. This holds true for all frequency bands except for gamma **(Figure 3-1E)**. All obtained single-session CAPs can be grouped based on their spatial similarity to obtain a basis set of CAPs that represent brain states that are present across mice (**Fig4A**). This holds for all signal types (not pictured), highlighting that obtained single-session CAPs, despite explaining only a part of the variance in the original data, represent distinct brain states that are consistent across sessions and mice.

To obtain the basis set of CAPs per frequency band, hierarchical clustering was performed on the CAP-CAP similarity matrix for each frequency band, such as the one shown in **Figure 4A**, using average linkage. Aggregated CAP groups were then defined by cutting the dendrogram at a fixed distance threshold (cutoff = 0.8), producing a set of spatially coherent CAP clusters shared across sessions. This cutoff was selected after visual inspection of the optimal leaf order of the resulting dendrogram across frequencies (**Fig4D**). The obtained basis set from this method was used to highlight the fact that CAP patterns are recurring across sessions and mice. Because the resulting CAP structure depends on the clustering cutoff, the basis set should be interpreted as a representative organization of the data rather than a definitive set of CAPs. All subsequent results were also obtained using cutoffs of 0.7 and 0.9. The obtained basis set of CAPs across frequencies, like the single session CAPs, represents spatial configurations that encompass previously characterized wide-field resting state networks (**Fig4-1A**). Key spatial configurations are also conserved across timescales and can broadly be represented by two anticorrelated CAP pairs (**Fig4-1B**).

**Figure 4.**
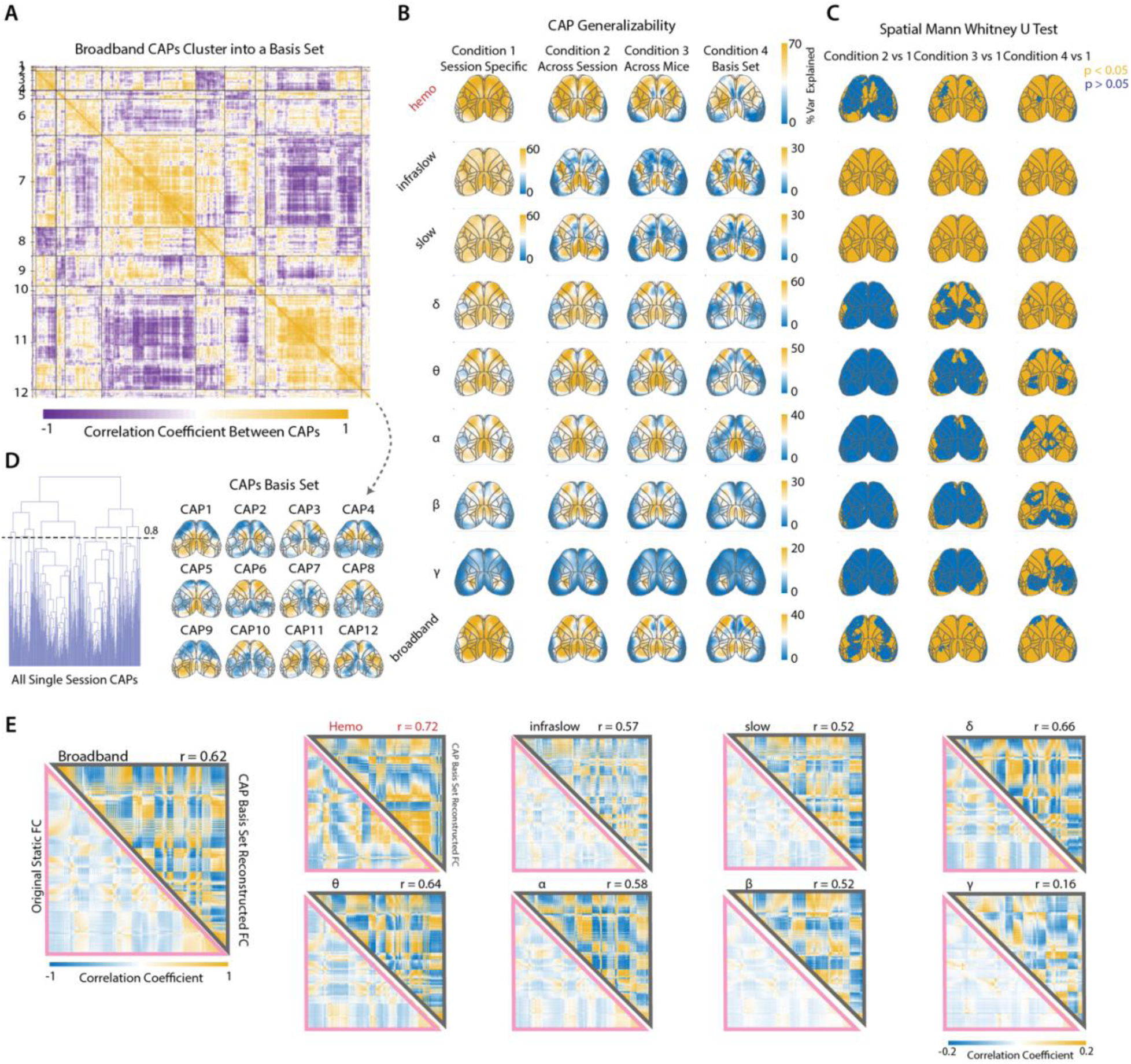
Spatiotemporal Generalizability of CAPs. **A)** The CAP-CAP spatial correlation matrix of all session-level CAPs for the indicated frequency band, reordered to group CAPs according to hierarchical clustering (average linkage; distance = 1 − spatial correlation) and cluster assignment at a distance cutoff of 0.8. Each pixel shows the spatial correlation between a pair of CAP maps. Blocks indicate sets of highly similar CAPs within the same aggregated cluster. Tick labels denote aggregated cluster IDs and black lines mark cluster boundaries. Below is the basis set obtained from clustering the broadband CAPs shown in the matrix. **B)** Maps denote the degree of variance that CAPs explain relative to the original data across all frequency bands. The first column represents condition 1 which is the average (n=28 sessions) degree of variance explained using the session-specific CAPs. Condition 2 shows the variance explained by reconstructing the data using specific CAPs obtained from the same mouse but from a different session. Condition 3 is then using CAPs from a different session and a different mouse. Condition 4 reconstructs the data using the basis set. Note the different colorbar for each row which is adjusted to scale based on the max explained variance in the session-specific map. **C)** Spatial map representing the statistically significant differences in explained variance between condition 1 relative to the other 3 conditions. Colors represent the FDR corrected Mann-Whitney U-Test thresholded p-values. Blue pixels have a corrected p-value>0.05, and yellow p<0.05. **D)** Dendrogram showing the clustering of all the broadband single session CAPs. 0.8 is the cutoff used for obtaining the basis set. **E)** Bottom left pink triangle represents the average FC matrix obtained from the raw data. This is the same data as in Figure 1G shown here to highlight a comparison to the top triangle. Top grey triangles represent the average matrix obtained from reconstructing session-level data based on the group-level basis set CAPs. The spatial correlation between the two matrices is displayed to the right of each square. Represented is the data for all frequency bands and signal types.

**Figure 5.**
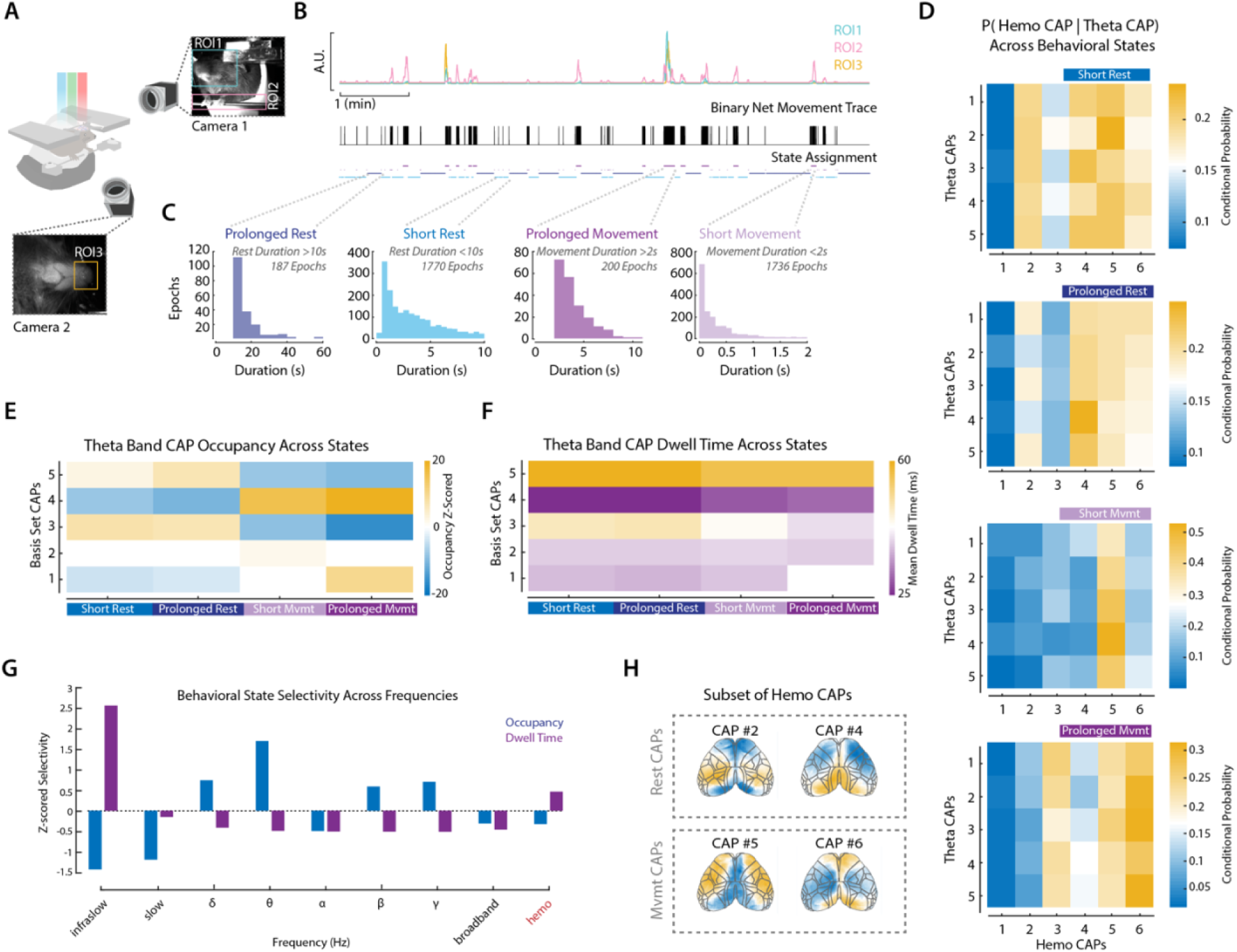
Behavioral State Modulates CAP Dynamics Through Frequency-Specific Mechanisms. **A)** Schematic illustration of camera set-up. FOV of camera 1 and 2 is shown as well as the area that encompasses each of the three ROIs used to assess motion. **B)** Top panel is an example session showing the time-varying change in motion energy extracted from the 3 ROIs shown in panel A. The middle panel is the net binary movement trace obtained from the combined movement information across all three ROIs. Based on the duration of a movement period, each frame is assigned to one of four different behavioral states: prolonged rest (>10s), short rest (<10s), prolonged movement (>2s) and short movement (<2s). **C)** Distribution across all sessions showing the number of epochs for each condition. **D)** Group-level occupancy across 4 behavioral states for the 5 basis set CAPs obtained for the theta band. Displayed is the z-scored difference between the observed probability of observing a CAP occurring in a given state and the probability expected under shuffled behavioral labels. Occupancies that is not statistically significant are set to zero (white squares). Significance was assessed using a two-sided permutation (empirical) test with FDR correction. **E)** Same as in panel D but showing the dwell time in ms for each CAP across states. **F)** Pairwise correlations between CAP rows in panel D. Repeated for all row pairs, across all the different frequency bands. Boxplots represent the distribution of these correlations. +1 means CAPs behave similarly across states while -1 means that they behave oppositely. **G)** Behavioral state selectivity across frequencies for both occupancy (blue) and dwell time (purple). >0 represents a higher-than-average selectivity for occupancy or dwell time. <0 means a lower-than-average selectivity relative to all other frequency bands.

To further assess the degree of generalizability of CAPs, the percent variance explained per pixel was calculated across all frequency bands (**Fig4B**). **Figure 4B** summarizes the generalizability of CAPs across four conditions: session-specific fitting (Condition 1), generalization across sessions within the same mouse (Condition 2), generalization across mice (Condition 3), and reconstruction using a group-level basis set (Condition 4). As expected, session-specific CAPs yielded the highest % variance explained across all frequency bands. Low-frequency bands (infraslow and slow) demonstrated relatively strong reconstruction that progressively declined when CAPs were generalized across sessions and mice. In contrast, higher-frequency bands (alpha–gamma) exhibited lower overall % variance explained across all conditions, but the magnitude of reduction between session-specific and generalized conditions was comparatively modest. Thus, while slower frequencies account for more variance overall, their reconstruction appears more sensitive across conditions.

The spatial structure of the explained variance is broadly conserved across conditions but is notably different across frequency bands (**Fig4B**). The variance explained in the hemodynamic and broadband voltage is broadly distributed across the cortex. In the band-limited data from delta to beta, the explained variance becomes gradually more localized to RSP and M2, both of which are key areas in the DMN. Gamma band variance explained exhibits a different spatial pattern, with most of the explained variance coming from V1 and M2. Pixel-wise Mann–Whitney U tests comparing session-specific reconstructions (Condition 1) to generalized reconstructions (Conditions 2–4) reveal a frequency-dependent pattern of statistical differences (**Fig4C**). Across-session (Condition 2 vs 1) and across-mouse (Condition 3 vs 1) comparisons show more widespread significant reductions (p < 0.05) in low-frequency bands, indicating that slow CAP structure is more sensitive to session and animal variability. In contrast, higher-frequency bands are dominated by non-significant pixels (p > 0.05), suggesting that CAP reconstructions at these frequencies remain statistically comparable to session-specific fits. The basis-set condition (Condition 4) exhibits broader significant differences across frequencies, reflecting the greater abstraction imposed by group-level clustering. Together, these results demonstrate that although low-frequency CAPs explain more variance overall, their generalizability is more sensitive across conditions, whereas higher-frequency CAP structure is comparatively stable across sessions and mice. These results also highlight frequency-dependent regional differences.

However, despite the reduced explained variance of low-frequency CAP basis sets, reconstructed data using the CAP basis set preserved the static FC structure with a correlation coefficient ≥ 0.52 for all frequencies except gamma (**Fig4E**). This indicates that the shared spatial scaffold of network co-activation is retained, while differences in variance primarily reflect session-specific temporal dynamics that do not substantially alter second-order correlation structure.

Taken together, these findings indicate that similar patterns of functional connectivity can emerge from distinct underlying network dynamics. Across frequencies, CAPs differed substantially in the amount of variance they explained, their persistence, and their generalizability, yet these differences had relatively little impact on the resulting static FC structure. This suggests that large-scale cortical networks are constrained by a common spatial architecture that is preserved across timescales, even as the temporal dynamics that generate these networks vary considerably. As a result, similar functional connectivity patterns can arise from different combinations of brain states and state transitions. The correspondence between neuronal and hemodynamic functional connectivity likely reflects this shared organizational framework, whereby slower hemodynamic signals capture the dominant spatial motifs of network activity while averaging over faster neural dynamics. More broadly, these findings further suggest that static FC alone may obscure important frequency-dependent differences in brain dynamics, motivating a direct examination of how brain states transition across neuronal and hemodynamic timescales.

### Behavior State Drives Slow Network Persistence and Fast Network Selection

The preservation of functional connectivity across frequencies suggests that there exists a common spatial scaffold of cortical organization. However, if similar connectivity patterns can emerge with distinct underlying dynamics, then time-varying changes in behavioral state may differentially influence how these network configurations are expressed across timescales. We therefore examined the relationship between behavioral state and CAP dynamics across neuronal and hemodynamic signals. Awake resting-state recordings are characterized by fluctuations in behavioral state occurring on the order of seconds [19], reflecting changes in arousal [23, 24] and often accompanied by subtle movements such as fidgeting [25]. These dynamic processes are known to modulate widespread patterns of neural [14] and hemodynamic activity [1] and are associated with specific CAPs [26], linking moment-to-moment behavioral variability to recurring large-scale brain states [15, 25, 27]. The behavioral state of the animal was classified into one of four categories (**Fig5A**). Rest periods are defined as either prolonged (> 10 s) or short (< 10 s). Movement periods are also broken down into prolonged movement (>2 s) and short (< 2 s) (**Fig5B** and **C**) since extended movement periods are generally associated with additional body movements [19] and have a distinct vascular response [28]. This provides us with a sensitive readout of movement periods, which includes gross orofacial movements (ROI1), forepaw movements (ROI2), and nose/whisking periods (ROI3). Across sessions, the most commonly occurring epochs were the short rest and short movement periods (**Fig5C**).

Taking the CAP basis set for each frequency band, which represents the group-level CAPs, we quantified the relationship between CAP expression and behavioral state by computing state-dependent CAP occupancy (**Fig5D, Fig5-1A**). CAP occupancy was defined as the conditional probability of observing each CAP given a behavioral state. To assess whether observed occupancy patterns exceeded chance levels, these values were compared to a shuffled control null distribution. The observed occupancy matrix was then z-scored relative to this null distribution, producing a z-scored occupancy (enrichment) matrix that reflects the degree to which each CAP is over- or under-represented in each behavioral state compared to chance (**Fig5D**). Statistical significance was assessed using a two-sided permutation test and corrected for multiple comparisons using FDR. Occupancy reflects the fraction of time that a given CAP occurs during a given behavioral state. There were certain CAPs that are preferentially expressed in rest versus movement periods. For example, CAP 3 and 5 were preferentially expressed during the rest periods, while CAP 4 is expressed in the movement states (**Fig5D**).

To test if the preferential occupancy is due to the fact that each CAP occurs many times within a given state or because it lasts a long time each time it occurs, we also calculated the dwell time across behavioral states. Dwell time was defined as the duration (in frames) of each CAP event. For each CAP and behavioral state, we computed the mean of dwell times across all events. This resulted in a CAP-by-state dwell time matrix for each frequency band (**Fig5E, Fig5-1B**), capturing how long each CAP persists under different behavioral conditions. In contrast to the occupancy, dwell time reflects how long a given CAP is maintained once engaged; for theta band CAPs, there is less of a distinction across behavioral states, but differences are larger across CAPs. Where CAP 5 has the longest dwell time across conditions, while CAP 4 has the lowest (**Fig5E**). This suggests that behavioral modulation in theta is driven more strongly by changes in CAP occurrence/selection than by prolonged CAP persistence.

To compare how behavioral state modulates CAP dynamics through changes in occurrence versus dwell time, we quantified state selectivity separately for occupancy and dwell time across frequency bands. For occupancy, state selectivity was defined using the z-scored occupancy matrix (CAP × state) by computing, for each CAP, the variance of the occupancy across behavioral states. These values were then averaged across all CAPs within a band to yield a single measure of occupancy-based state selectivity per frequency band. These values are plotted as the Z-scored selectivity across frequency bands by normalizing values to the mean and standard deviation of the occupancy-based selectivity values across frequency bands (**Fig5G, blue bar plots)**. Dwell-time selectivity was computed using the event-based dwell time matrix (CAP × state), where dwell time reflects the average duration of contiguous CAP events within each behavioral state. To assess the degree of spread of dwell times across states, the range (maximum minus minimum dwell time across a CAP) was calculated and then averaged across CAPs to obtain a band-level measure of dwell-based state selectivity (**Fig5G, purple bar plots)**. Similarly to the occupancy-based selectivity, these values were normalized across frequencies to be plotted side-by-side across frequency bands, enabling direct comparison of how strongly behavioral state modulates CAP occurrence versus persistence (**Fig5G**).

This analysis revealed that infraslow and hemodynamic CAPs are more modulated by behavior through changes in dwell time as represented by a higher difference in dwell time across behavioral states compared to other frequency bands (**Fig5G, Fig5-1A and B**). In contrast theta band reflects state-dependent selection of CAPs rather than stability (**Fig5G**), as represented by a higher variance in occupancy across behavioral states compared to other frequency bands. CAP dwell times in slow-gamma bands are also less differentiated across behavioral states compared to the other frequency bands (**Fig5G)**. These findings were robust to using different cut-off values for the CAPs basis set (tested also with cluster cutoffs of 0.7 and 0.9) (**Fig5-2C, Fig5-3C**). Other trends, such as gamma-bands state-dependent selection of CAPs rather than stability, were not consistently observed across CAP cutoffs.

These findings demonstrate that behavioral state influences large-scale cortical dynamics through distinct mechanisms across timescales. Thus, similar behavioral transitions can be represented either by stabilizing ongoing network configurations or by altering which configurations are preferentially expressed, depending on the temporal scale of the underlying activity. These results provide evidence that the conserved functional connectivity architecture identified in earlier analyses can arise from fundamentally different dynamic processes. More broadly, they suggest that slow activity may reflect sustained changes in global brain state, whereas faster activity supports more flexible reconfiguration of network structure in response to moment-to-moment behavioral demands.

## DISCUSSION

In this work, we used simultaneous wide-field voltage and hemodynamic imaging in awake mice to demonstrate that hemodynamic networks preserve the spatial organization of large-scale cortical activity but capture only a subset of its frequency-dependent dynamics. Across neural frequencies spanning infraslow to gamma, canonical functional networks maintained a largely conserved spatial organization, despite systematic differences in connectivity strength and temporal expression. Importantly, we found that behavioral state modulated these networks through distinct temporal mechanisms: slower neural dynamics were primarily shaped by changes in network persistence, whereas faster dynamics were characterized by changes in network occurrence. Hemodynamic network dynamics closely resembled those of infraslow neural activity, exhibiting similar behavior-dependent changes in state persistence but failing to reflect the faster changes in network occurrence. Together, these findings demonstrate that hemodynamic functional networks primarily capture the persistence of slow neural network states rather than the full range of behavior-dependent neural dynamics, providing a framework for interpreting the neural processes underlying resting-state hemodynamic connectivity.

### Functional Connectivity Emerges from Layered Neural Dynamics Operating at Distinct Temporal Scales

FC captures coherent correlations in spontaneous activity between distant brain regions and is most commonly used to analyze resting-state fMRI data. These correlated fluctuations in the hemodynamic signal are typically interpreted as reflecting coordinated neuronal population activity underlying functional brain networks [29]. However, similar large-scale network organization is not unique to slow hemodynamic signals and has also been observed in neuronal activity across a broad range of timescales using modalities such as wide-field optical imaging [15, 30], EEG [5–7], and electrophysiology [31, 32], indicating that large-scale functional network structure is a fundamental property regardless of brain dynamics. Further, this network structure is independent of signal modality (hemodynamics vs. voltage). Our Results extend this picture by demonstrating that canonical functional networks are preserved in time-averaged FC across all neuronal frequency bands from infraslow to gamma. The repertoire of available spatial configurations is largely invariant across timescales. Despite this shared scaffold, the strength of connectivity systematically decreased with increasing frequency, consistent with reduced temporal persistence of coordinated activity. Notably, modality-specific differences emerged: the hemodynamic signal exhibited stronger connectivity within somatosensory regions, whereas the voltage (JEDI) signal showed enhanced correlations within the default mode network (DMN) and between DMN and other networks. In contrast, we observe no meaningful differences between mCherry and dR/R signals, suggesting that the mCherry signal captures the same underlying hemodynamic functional organization as the dR/R signals.

A growing body of work suggests that time-varying brain activity can be described using a lower-dimensional set of recurring patterns, a phenomenon consistently observed across fMRI [9, 21], wide-field imaging [4, 12], and large-scale electrophysiological recordings [33]. Our CAP analysis extends this framework in a critical direction: we show that the difference in FC strength across frequencies arises not from fundamental differences in the spatial organization of networks, but from the differences in the temporal dynamics with which a common set of network configurations can be expressed. At low frequencies, CAP dynamics were dominated by prolonged dwell times, producing sustained correlations and stronger FC. In contrast, higher-frequency CAPs that preserve similar spatial organization were more transient and rapidly recurring. These CAPs contributed less to long-timescale correlations. Static time-averaged FC therefore primarily reflected the temporal persistence with which a shared set of large-scale network configurations was expressed, not the spatial architecture of those configurations itself. This implies that because the spatial scaffold is held constant across frequencies, differences in static FC magnitude can be attributed specifically to differences in temporal dynamics, and static FC alone cannot resolve those dynamics.

### Behavior Shapes the Temporal Organization of Large-Scale Cortical Activity

A growing body of work suggests that spontaneous brain activity reflects a low-dimensional dynamical system in which spatially structured fluctuations give rise to coordinated large-scale networks. These dynamics are closely linked to behavioral and arousal states, particularly during quiet rest, where strongly fluctuating patterns of activity across distributed regions contribute substantially to interregional correlations [1]. Recent studies further showed that spontaneous behaviors are accompanied by rapid changes in both the magnitude and correlational structure of cortical activity, underscoring the tight coupling between behavior and large-scale brain dynamics[15, 34]. Our results build on this framework but reveal a dimension of behavioral modulation that has not previously been characterized: the mechanism through which behavior reshapes network dynamics is itself frequency-dependent. At low frequencies for both infraslow neural and hemodynamic activity, behavioral modulation primarily altered the persistence of network states, leading to prolonged dwell times and sustained periods of coordinated activity in either prolonged movement or prolonged rest periods. In contrast, higher-frequency activity was dominated by changes in state occurrence, with stable network motifs being flexibly recruited over time rather than persistently maintained. The same behavioral transition, from rest to movement, was therefore represented through opposite temporal mechanisms depending on the frequency band examined.

The persistence-versus-selection dissociation is the central mechanistic finding of this work and has several important implications. First, it demonstrates that static functional connectivity is mechanistically ambiguous. Identical FC patterns can be generated by prolonged state persistence at one timescale or by differential state selection at another, meaning that changes in FC observed across behavioral conditions or clinical populations cannot be interpreted as reflecting a single underlying process without frequency-resolved dynamic analysis. Despite differences in dynamic expression across frequencies, reconstructed activity preserved the overall static FC structure, even when low-frequency CAP basis sets explained less variance across sessions than session-specific CAPs. This suggests that large-scale functional organization is constrained by a stable spatial scaffold, whereas behavior primarily modulates the temporal expression and sampling of these network configurations across timescales.

### Hemodynamic Network Dynamics Reflect Slow Persistent Neural Networks

Our simultaneous voltage and hemodynamic measurements further clarified what aspects of neural activity can be captured by vascular signals. While hemodynamic FC preserved canonical network structure, consistent with a shared spatial scaffold, its dynamics were strongly dominated by low-frequency components and exhibited pronounced dwell-time modulation, just like the infraslow voltage signal. This finding directly addresses a limitation of prior hemodynamic only resting-state studies in mice, which could characterize the spatiotemporal structure of fMRI CAPs but could not determine which temporal regime of neural activity those patterns reflect. Critically, hemodynamic CAP dynamics are most closely paralleled by infraslow neural activity specifically; both exhibited pronounced dwell-time modulation by behavioral state, with behavior primarily acting through sustained changes in network persistence rather than rapid shifts in state selection. Higher-frequency neural dynamics, characterized by rapid state transitions and behavior-driven selection, were not reflected in the hemodynamic signal. Thus, hemodynamics provides a behaviorally modulated, low-pass filtered representation of neural dynamics that preserves large-scale spatial organization while compressing temporal variability.

These findings provide a mechanistic interpretation of fMRI resting-state measures, which primarily index the persistence of slow network states, not the full repertoire of dynamic network activity. This has direct implications for interpreting resting-state fMRI. Behavioral or arousal-related changes in fMRI connectivity are therefore most likely to reflect changes in how long slow network configurations are maintained, not changes in which configurations are selected or how rapidly they transition. Conditions that alter fast network dynamics without changing slow persistence may be invisible to hemodynamic measures, while conditions that stabilize or destabilize slow network states will produce the largest hemodynamic FC effects. Simultaneous voltage and hemodynamic imaging of the kind demonstrated here provides a means to directly dissociate these contributions and ground hemodynamic connectivity measures in their neural basis.

Our findings have some limitations that can be addressed with future investigations. First, our trial durations are relatively short, constrained by the high sampling rate and limits on the number of frames that can be stored per trial with our imaging system. This limitation is particularly relevant for lower-frequency analyses, where longer recordings may be necessary to fully capture slow dynamics and a broader range of behavioral variability. As such, low-frequency CAPs may exhibit greater convergence across sessions and explain more variance if longer trials were used to better sample diverse brain and behavioral states. Second, while the mCherry signal was used both for hemodynamic correction and as a proxy for hemodynamic signal changes, it provides a less direct readout compared to approaches that use interleaved diffuse reflectance imaging of red and green light [20]. Although similar strategies have been successfully employed in studies, the mCherry signal remains a composite optical measure of hemoglobin absorption and fluorescence, similar in many ways to the hemodynamic signal obtained from the ratiometric VSFP-Butterfly sensor. While also not a perfect readout of HbT, HbR or HbO, VSFP-Butterfly has proven to be useful in many studies relating neural to hemodynamic activity [27, 35, 36]. We believe mCherry can serve a similar role when performing voltage imaging at much higher frame rates, where interleaving of additional LEDs for hemodynamic correction are not feasible. Thirdly, we derived CAPs from band-limited signals, and temporal filtering may introduce dependencies that transform discrete state transitions into continuous mixtures of underlying patterns. As a result, the observed variability in low-frequency CAPs reflects not only neural dynamics but also the interaction between temporal integration and behavioral variability. Clustering approaches impose a discrete representation onto what is likely a continuous state space, and the number of CAPs may differentially constrain single-session and group-level analyses, particularly at low frequencies where greater heterogeneity may require higher model complexity.

One possible interpretation of our findings is that cortical dynamics reflect at least two partially overlapping modes of state organization operating on different timescales. Slow resting state fluctuations, which dominate hemodynamic and low-frequency voltage activity, may reflect gradual global changes in behavioral state or internal state that stabilize network configurations over extended periods. In contrast, faster neural activity may capture more rapid transitions associated with moment-to-moment behavioral engagement, enabling flexible selection of network configurations without prolonged persistence. Under this framework, slow activity shapes the background state in which cortical networks operate, whereas faster dynamics support rapid reconfiguration within that state. Rather than reflecting distinct network architectures, these dynamics may represent complementary temporal modes through which the same large-scale cortical networks are organized and expressed.

## METHODS

### Animals

4 female and 1 Male *EMX1-Cre* mice were used for experiments. Mice were housed in a reverse cycle (12h light, 12h dark). All experiments were performed during the dark cycle. Mice were provided with *ad libitum* food and water. All mice had a running wheel as enrichment in their cages and were either singly housed or with their same sex litter mates. Emory University Institutional Animal Care and Use Committee approved all animal work performed in this study.

### Neonatal intracerebroventricular injection

Detailed methods can be found in a previously published article [16]. Briefly, to achieve brain-wide expression of green JEDI-1P and red mCherry we used intracerebroventricular injections in neonatal pups to deliver a mixture of two viral vectors AAV.PHP.eB-EF1a-DIO-JEDI-1P-Kv2.1-WPRE (2–4×10^12^ vg/mL) and AAV9-hSyn-mCherry (1.3 × 10^13^ vg/mL). These vectors were mixed with a particle ratio of 1:1.3. A total of 2uL of the viral vector mixture was delivered to each ventricle via a 10uL Nanofil syringe (World Precision Instrument) with a 34G beveled needle (NF34BV-2, World Precision Instrument).

### Imaging Set-up

For through-skull imaging mice were implanted with a clear-skull window and headpiece [37]. Fluorescent signals were imaged from awake, head-fixed mice using a macroscope (MiCAM Ultima, Brainvision). The imaging plane of the macroscope was focused through the intact skull slightly below (∼100 µm) the most superficial blood vessels. Both JEDI-1P-Kv and mCherry were excited with a single light provided by a blue LED (LEX2-B, Brainvision) conditioned through a bandpass filter (FF01-466/40, Semrock) and a dichroic mirror (FF495-Di03, Semrock). Emitted green JEDI-1P and red mCherry reference fluorescent signals were split by a second dichroic mirror (FF580-Di01, Semrock) and recorded simultaneously using two CMOS cameras (100 × 100 pixel array, 100 × 100 µm per pixel). The first camera with emission filter (FF01-525/50-50, Semrock) recorded JEDI-1P-Kv while the second one with emission filter (FF01-650/60-50, Semrock) recorded mCherry fluorescent signal. The camera acquisition was controlled by the MiCAM acquisition software. Acquisition frame rates for both cameras were set to either 40 or 200 Hz. The power of the blue excitation LED was 3.86 mW (intensity 0.05mW/mm^2^) at the plane of the imaging window for all experiments.

Red reflectance imaging was done using the same set-up described above, including all the dichroic and emission filters. However, only the signal from one camera with the 650 nm emission filter was used. The light source was a white light conditioned with a red filter (FF01-632/22-55nm, Semrock). The power in the plane of the imaging window for the diffuse reflectance experiments was 291 µW (intensity 4 µW/mm^2^).

### Image Acquisition Paradigm

Each imaging session consisted of two different types of recording sessions: whisker stimulation and resting state trials. Some sessions also differed in terms of the frame rate used (40Hz vs 200Hz) and the LED light source (blue vs red). The data acquisition will be described in the order that it is displayed in **Supp Fig 1D**. Note that across session days, the order of red versus blue excitation light was switched to control for possible slow changes in general arousal levels that could occur throughout the duration of an imaging session.

For fluorescence imaging, the blue LED was used. The first set of trials consisted of whisker stimulation trials that were 10.24s in length and contained 3s long 30Hz whisker air puffs (50% duty cycle) that were delivered to either the left or right side of the mouse’s face. This data was acquired at 200 Hz. Several resting-state trials were then acquired using either a frame rate of 200Hz or 40Hz. Our imaging system saves frames in between trials and, therefore, has a limit on the number of frames that can be stored for a given trial. This limit influences the maximum trial duration of a trial which can be modulated by the selected frame rate. We set up trials to acquire the maximum number of allowable frames for each trial type. The 200 Hz trials are used for the broadband signal analysis looking at infraslow to gamma band activity. Each trial acquired at this frame rate is 40.96s long.

After fluorescent imaging, the excitation light source was switched to acquire diffuse red reflectance data in the same mice. Here we again obtained both whisker stimulation and resting state data. The final resting state dataset used for analysis for each mouse consists of 102.3 min (at 40Hz) and 40.98 min (at 200Hz) of JEDI fluorescence imaging and an additional 102.3 min (at 40Hz) of red reflectance imaging.

The final dataset of JEDI-1P imaging consisted of a total of 289 resting state trials at 200FPS and 540 whisker stimulation trials. This data was collected from 5 mice, with a total of 3 – 7 recording sessions. Red reflectance data is collected from 4 mice, and consists of 120 resting state trials at 40FPS, and 562 whisker stimulation trials.

### Image Preprocessing

The preprocessing pipeline for fluorescence imaging is a modified version of the pipeline previously designed and published by our group [16] and is also outlined schematically in **Supp Fig1E**. All image registration is done using rigid transforms, which includes only translation and rotation. Imaging frames acquired from both cameras are first registered to the trial averaged image across all frames obtained from Camera B (the green color channel). Across-trial registration was then done by registering all trials to the time-averaged image obtained from the first trial. Both were done using the MATLAB imregtform function. Background subtraction was applied by subtracting the ‘dark’ image acquired for each mouse per session. This is a single 10.24s long trial acquired for each mouse per recording session without any excitation light. The trial average fluorescence obtained is subtracted from all data for that session to account for the dark noise of the camera and changes in ambient light conditions. Next, we account for photobleaching in both color channels by fitting a bi-exponential model (per pixel) to the trial-averaged data for a given session type. Using this fitted model, single trials are detrended. Change in fluorescence was then calculated by taking F0 to be the mean of the full trial for resting state trials and the mean of the first second before stimulation onset for the whisker stimulus condition. Then, for each session, a manual mask was drawn to conservatively mask out all pixels that are outside of the imaging window as well as pixels that contain noticeable air bubbles.

Hemodynamic and motion artifact contamination of the green JEDI-1P-Kv signal was removed using filtered versions of the simultaneously acquired ΔF/F0 red reference channel. A sequential regression step is used to scale and subtract filtered versions of the mCherry trace from the JEDI-1P-Kv signal using Ordinary Least Squares Regression, as outlined in more detail here [16]. For trials sampled at 200Hz, filters with cutoffs at [14–30], [10–14], [5–10], and [1–5] were applied to the reference channel. Note that for trials sampled at 40Hz, the [14-30Hz] filter was omitted. Slower hemodynamic changes were removed from the JEDI-1P-Kv trace using the <1Hz global signal regressed (GSR) mCherry. This slow <1Hz GSR mCherry trace was also used as our measure of hemodynamics. The JEDI-1P-Kv ΔF/F0 was lowpass filtered at 70Hz, and then using sequential regression, filtered versions of the red reference channel as described were removed from the JEDI-1P-Kv signal. This preprocessed JEDI-1P-Kv trace is the presented measure of changes in membrane voltage potential used throughout the paper unless stated otherwise.

Red reflectance imaging trials were preprocessed similarly using only a subset of steps. These included image alignment across trials, background subtraction, manual masking, and ΔF/F0. This was done as described above (**Supp Fig1F**).

Group-level analysis was done by aligning all session data to the Allen Atlas. Whisker barrels in each session were identified by looking at increases in power of the JEDI-1P-Kv signal at 30Hz (which corresponds to the whisker stimulus frequency). By plotting the average difference in power between the stimulation period and baseline, a map with clear activity in left and right whisker barrel is obtained (**Supp Fig1G**). Manual circular ROIs (radius =3) were selected for the left and right hemispheres that were centered on the pixels with a clear response. The centroid of these selected ROIs, along with the centroid of the Allen Atlas mask of barrel cortex was obtained using MATLB regionprops(). The transform between these two pairs of centroids was then calculated using a rigid transform with MATLB fitgeotform2d().

#### Fluorescent Signals Modeled from First Principles

##### The transmission of light in a scattering medium like the brain can be approximated through the modified Beer-Lambert Law

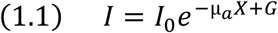

where *I0* represents the incident intensity, and *I* is the resulting intensity after the light has traveled a path length *X* through a medium that has µa absorption coefficient*. G* represents the geometric factors that introduce uncertainty regarding how much light is exiting the brain as detected by the camera. Measurements of intensity are often expressed in relative terms (by dividing by I (t=0)) which removes influences of *I0* and *G* which are challenging to calibrate. By incorporating wavelength and time dependencies into equation (1.1) we get

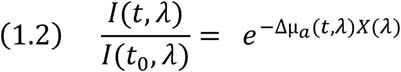

While *Δµa* is represented as

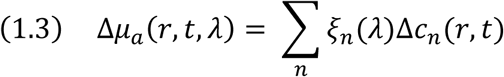

Here *µa* is a function of *r* which denotes a given location in 2D space, for a given time point *t* and wavelength *λ*. *ξn (λ)* refers to the extinction coefficient of a given absorber while *Δcn(r,t)* is the change in concentration of a given absorber relative to an earlier time point. In the visible range (400-700nm), the major time-varying absorbers of the brain are HbO and HbR. Therefore, *Δµa* for wide-field optical imaging takes into consideration changes in the concentration of HbO and HbR. The wavelength-dependent path length and the absorption coefficients for both absorbers in the brain have been modeled[20].

Taken together equation (1.2) can be used express changes in fluorescence of both JEDI-1P-Kv and the red reference channel mCherry (**Supp Fig2A**). For JEDI-1P-Kv the change in measured fluorescence is a function of both the excitation wavelength (466nm) and the emission wavelength (525nm). Note from here on we use 526nm rather than 525nm for the modeling, because the closest published absorption values for HbO/HbR are available for 526nm [20].

The change in intensity can be represented as:

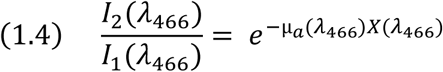

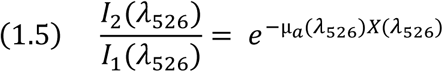

In the case of a voltage indicator, the change in fluorescence is not only a function of changes in the concentration of absorbers but also varies as a function of changes in voltage membrane potential expressed as *Cf(t)*. Combining (1.4) and (1.5) the observed change in fluorescence of JEDI-1P-Kv is thus represented as

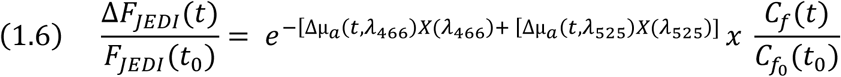

A similar equation can be derived for the red reference channel mCherry which has the same excitation wavelength (466nm) but a different emission wavelength (650nm).

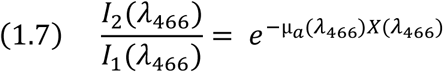

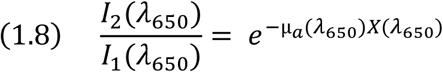

Where the measured change in fluorescence can be denoted as a combination of (1.7) and (1.8)

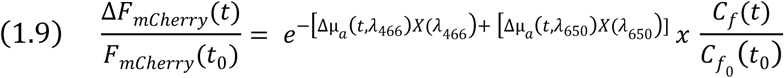

Note that the fluorescence of mCherry does not change as a function of time, setting the last term in (1.9) equal to 1. Thus, the measured change in fluorescence of the mCherry (1.9) is only a function of changes in concentration and subsequent absorption of HbO and HbR.

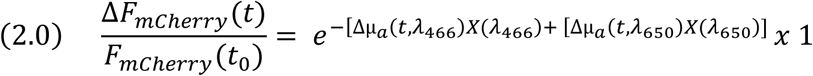

Using equations (1.6) and (2.0) we can model how the measured change in fluorescence is expected to vary in response to a whisker air puff stimulus which through neurovascular coupling elicits changes in concentration of HbO/HbR. To do this, we used values of changes in concentration of HbO, HbR and HbT in response to a whisker air puff stimulus as reported by Ma and colleagues (**Supp Fig2B**) [20]. We also used reported values for changes in the extinction coefficient *ξn (λ)* for HbO and HbR, as well as the path length of light traveled at each given wavelength (**Supp Fig2C**) [20]. Combining this information, we can model expected changes in light absorption for excitation, emission, and the net recorded intensity for a whisker stimulus. JEDI-1P-Kv is most sensitive to changes in HbT for both the excitation and emission wavelengths of light. This is evident by the decrease in net light intensity following onset of the whisker stimulus (**Supp Fig2D, left panel**). The red reference channel in contrast is a combination of HbT sensitivity driven by the excitation wavelength and HbR driven by the emission wavelength. Due to the significant order of magnitude difference however between the absorption coefficients at shorter wavelengths of light the net change in light intensity also trends in a negative direction, suggesting that this signal is more weighted towards HbT (**Supp Fig2D, right panel**). Taken together the slow hemodynamic contaminating in both signals results in a net decrease in light intensity for both channels (**Supp Fig2E**) which mirrors the results we see *in-vivo* (**Fig1C**).

### Behavior Monitoring

Spontaneous changes in behavior of the animal was monitored continuously throughout each session using 2 Basler cameras imaging at 100FPS (acA1440-220um, Basler Camera). The first was positioned to the side of the mouse, and captured forelimb paw movements and gross orofacial movements. The second camera was positioned directly below the animal and captured their full bottom view of their face from nose to neck.

Spontaneous changes in behavior were obtained by calculating the squared temporal differences of mean frame-to-frame changes in signal intensity for 3 different regions of interest (ROI)[27]. These three ROIs include the area spanning the bars where the paws rest, the side of the face and the nose tip (**Fig4A**). Frame-wise displacement was calculated for each ROI across all sessions. For each ROI per session, changes in frame wise displacement that are 1 STD away from the mean are considered to be movement periods. All behavior metrics are very tightly linked. Orofacial movements from the side of the face and nose movements are most strongly linked (cross- correlation average value of 0.7 with a time lag of 0.06s). While forelimb movements are slightly less correlated and follow the onset of face/nose movements (cross-correlation average of 0.45, time lag -0.11s to nose and -0.13s to face).

To have a conservative estimate of behavioral state that includes any type of movement the animal makes within any of the three ROIs, a net movement trace for each session is obtained by combining the movement information from each ROI. To mirror behavioral state analysis done in prior studies each behavioral trace was then split up into short (<2s) and long (>2s) movement bouts [19]. Rest was also split into prolonged rest (>10s) and transition periods of rest between movement bouts (<10s). Behavioral data was simultaneously recorded for 25 sessions across 5 mice. This resulted in a group total of 1736 movement epochs, with 200 of these being longer than 2s. There are an additional 1770 rest epochs, of which 187 are longer than 10s.

### Bandlimited Signals

Throughout the paper, the <1Hz mCherry signal is compared to neuronal activity by looking at the relationship to bandlimited activity of the latter. For this changes in the bandlimited JEDI-1P-Kv signal were calculated. For ROI-based cross-correlation analysis the JEDI signal is bandpass filtered into smaller ranges with a frequency resolution of 2 Hz. For cortex-wide analysis the signal is filtered into the canonical bands of [0.05 – 0.1], [0.1 – 1], [1–4], [4–8], [8–13], [13–30], and [30–70] Hz. All bandpass filtering was done using a 4^th^-order bandpass FIR filter. Power modulation was then calculated by squaring the bandpass filtered signal and low pass filtering it using a 4^th^ order lowpass FIR filter with a cut-off of 1Hz for all frequency bands except those below 1Hz i.e. for [0.05 – 0.1] and [0.1 – 1] where the bandpass filtered signal was used instead. Most results in the paper rely on the bandpass-filtered signal rather than the power-modulated signal to preserve the anticorrelation between networks.

### Static Functional Connectivity

All imaging data is aligned to a common Allen Atlas space as described in our preprocessing pipeline. Static FC, both the seed based and the FC matrices, were obtained using the GSR data. Static seed-based connectivity maps were generated by correlating the average time varying signal from an ROI ranging between 26-28 pixels to all other pixels. To generate FC matrices the imaging data was not spatially smoothed or parcellated. Therefore, each FC matrix is [3617,3617] pixels in size each representing 100um by 100um. Across dorsal cortex there are 3 common brain networks that can be resolved: the default mode network (DMN), the somato-motor network (SoMo) and visual network (VIS). To facilitate comparison to other fMRI studies, FC matrices were generated by grouping pixels into these 3 networks. Due to importance of the DMN for resting state studies, all pixels from a given Allen Atlas regions that correspond to areas affiliated with the DMN were included in this network. All remaining areas were then subdivided into SoMo and VIS. FC matrixes were generated so that activity within each network from both hemispheres is plotted, as opposed to plotting all left hemisphere data from all networks and then all right hemisphere data from all networks. Note that the choice of how to arrange each ROI in the FC matrix is purely for ease of visualization. It has no effect on the results comparing the neuronal and hemodynamic derived FC. This also has no implication on further dynamic analysis which also does not rely on predefined parcellations or networks.

#### Static Functional Connectivity Statistics

For statistical significance testing all single-subjection correlation maps and matrices were Fisher R-to-Z transformed, averaged across animals and then thresholded to significant connections using a two-tailed t-test with p< 0.05, FDR corrected. Averaged maps were then re-transformed to correlation values (r-scores) for plotting. Comparisons between groups were done using the same procedure by averaging Z-transformed correlation values across mice within a group and comparing groups with a two-sample two-tailed, t-test with FDR correction.

### Session Specific Cortex Co-Activation Patters (CAPs)

To investigate the resting state dynamic functional organization, we extracted co-activation patterns (CAPs) for each session, frequency band and signal type. Each session consisted of 10 resting state trials. This method classifies wide-field imaging frames into clusters based on their spatial similarity, which has been done extensively using resting state fMRI data in human fMRI [21] and has since been used also in preclinical imaging [22]. The averaged frames within each cluster were taken to represent recurrent patterns of resting-state wide-field CAPs. The same preprocessed and aligned imaging data used for the static FC analysis was used here as well. Data for a given frequency band was concatenated across trials for a given session. CAPs for each session were then identified by running k-means clustering on the z-scored session level data (ran for 500 iterations, with 15 initializations, using KMeans function from Scikit-learn in Python). This clustering was done on a per session basis for clusters ranging in size from k=2 – 26. For each k we calculated the explained variance as well as the gain in explain variance with increasing cluster number. Explained variance here is defined as:

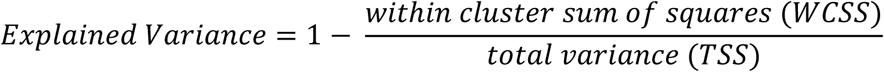

The ideal number of K clusters for each session was determined by identifying the elbow point, where the gain in explained variance levels off. This was done by finding the point of maximum curvature by identifying the peak of the second derivative.

### CAP Clustering for Basis Set

To identify group-level CAPs, we used an unsupervised hierarchical clustering algorithm. Group-level clustering was done on a per-frequency-band basis. For each band, we constructed a common basis set of CAPs by aggregating and re-clustering session-specific CAPs across animals and recording sessions. Here we take the ideal number of clusters for each session as determined using an elbow criterion described above. To avoid including rare or unstable patterns, clusters with occupancy below 2% of frames were excluded (this represents CAPs that were present for less than 7.2s across a resting state session) when the subset-optimized elbow solution differed from the full elbow estimate.

All remaining session-level CAPs within a given frequency band were pooled across sessions and animals. Each CAP was represented as a flattened spatial map (100 × 100 pixels) and z-scored across pixels to normalize spatial variance. Pairwise spatial similarity between CAPs was quantified using Pearson correlation, yielding a CAP–CAP correlation matrix. This similarity matrix was converted to a distance matrix defined as 1-r, where r denotes the spatial correlation. Hierarchical clustering was performed on this distance matrix using average linkage. Aggregated CAP groups were then defined by cutting the dendrogram at a fixed distance threshold (cutoff = 0.8), producing a set of spatially coherent CAP clusters shared across sessions. This cutoff was selected after visual inspection of the optimal leaf order of the resulting dendrogram across frequencies. The obtained basis set from this method was used to highlight the fact that CAP patterns are recurring across sessions and mice, but it does not represent a final “true” set of CAPs. Rather, it just suggests that activity can be represented by a subset of spatial patterns. Changing the cut-off results in more complex CAPs that further subdivide the networks observed here.

All analyses that relate CAPs to the behavioral state of the animal were run on three different cutoff values: 0.7, 0.8 and 0.9. With increasing cut-offs resulting in fewer selected clusters and thus a reduction in the ideal CAPs that are maintained for each basis set.

### Reconstructing Data with CAPs

To compare how well individual per-session CAPs generalize across sessions within a given mouse, CAPs from different sessions were used to reconstruct data. The ideal number of K CAPs from a different session was selected. To fit these CAPs to each time point from a different session, the squared Euclidean distance between each imaging frame and each CAP centroid was calculated. CAPs were assigned to the nearest frames, exactly as in standard k-means. However, instead of reconstructing the frame as the raw cluster centroid, we computed a least-squares scalar gain for that frame, effectively fitting the selected CAP to the data via a single multiplicative amplitude term to account for global differences across sessions and mice.

### CAP Occupancy and Dwell Time Across Behavioral States

CAP occupancy was defined as the conditional probability of observing each CAP given a behavioral state. This was calculated by normalizing CAP counts within each state by the total number of timepoints in that state. To assess whether observed occupancy patterns exceeded chance levels, we generated a null distribution by shuffling behavioral state labels within each session (using circular shifts to preserve temporal structure) across 1000 iterations. For each shuffle, occupancy was recomputed, yielding a null distribution of values. The observed occupancy matrix was then z-scored relative to this null distribution, producing a z-scored occupancy (enrichment) matrix that reflects the degree to which each CAP is over- or under-represented in each behavioral state compared to chance. Statistical significance was assessed using a two-sided permutation test, where p-values were computed based on the deviation of observed values from the null mean. These p-values were subsequently corrected for multiple comparisons across CAPs and states using the Benjamini–Hochberg false discovery rate (FDR) procedure, yielding q-values for each CAP–state pair. CAP temporal dynamics were assessed using an event-based dwell time analysis. Contiguous sequences of identical CAP labels were identified as discrete events within each session, and each event was assigned a behavioral state based on the modal state during that period. Dwell time was defined as the duration (in frames) of each CAP event. This resulted in a CAP-by-state dwell time matrix for each frequency band, capturing how long each CAP persists under different behavioral conditions.

## Acknowledgements

Many thanks to Dr. Yunmiao Wang for providing the initial version of the preprocessing code and for teaching me the neonatal injection method. Thanks to Francois St.Pierre and his lab for providing the JEDI-1P constructs.

## Funding

F31 NS134314-03 (L.M.B.)

R01 NS111470 (D.J.)

R01 NS140219, R21 MH140191, R01 NS078095 and R01 EB029857 (S.K.)

## Competing interests

The authors declare they have no competing interests.

## Data, code, and materials availability

All data will be made publicly available on DANDI in NWB format.

Code is available via the following link DOI: 10.5281/zenodo.21862174

## Notes

### Competing Interest Statement

The authors have declared no competing interest.

### Summary of Updates

The title, abstract, and discussion have been updated to highlight the key findings better.

